# Granulocyte-macrophage colony stimulating factor targets lung stem cell niches to accelerate alveolar repair after virus-induced lung injury

**DOI:** 10.64898/2026.08.25.746956

**Authors:** Ana Ivonne Vazquez-Armendariz, Theresa M. Schäfer, Learta Pervizaj-Oruqaj, Ali Khadim, Ioannis Alexopoulos, Monika Heiner, Ludmila Sperling, Katharina Humpert, Maximiliano R. Ferrero, Benjamin Ott, István Vadász, Rory E. Morty, Thorsten Hain, Elie El Agha, Susanne Herold

## Abstract

Influenza virus pneumonia causes severe damage of the lung parenchyma, resulting in respiratory failure. Timely and coordinated epithelial tissue repair is crucial for re-establishment of gas exchange. We identify granulocyte-macrophage colony-stimulating factor (GM-CSF) as a niche-derived growth factor produced in response to viral lung injury by distal epithelial progenitor cell populations, including alveolar epithelial type II cells (AECII) and bronchioalveolar stem cells (BASCs). Using complementary *in vivo* infection models, loss- and gain-of-function approaches, and lung organoid systems, we reveal that GM-CSF directly promotes distal epithelial progenitor cell expansion and alveolarization. Mechanistically, GM-CSF suppresses AMP-activated protein kinase activation and enables mechanistic target of rapamycin complex 1 (mTORC1) signaling, driving epithelial progenitor cell proliferation. Administration of recombinant GM-CSF during the initial days of infection enhances AECII proliferation and differentiation into AEC type I, accelerating lung barrier repair. Together, our findings establish GM-CSF as a key regulator of distal lung progenitor cell niches that couples cytokine signaling to metabolic control of tissue regeneration. These results uncover a previously unrecognized epithelial-intrinsic function of GM-CSF and highlight its therapeutic potential to promote lung repair in acute injury.

## Introduction

Acute respiratory distress syndrome (ARDS) is a severe complication of influenza A virus (IAV) infection characterized by damage of the lung epithelial barrier resulting in exudation of edema fluid into the alveolar compartment and impaired gas exchange^1^. The replenishment of alveolar epithelial cells type 1 (AECIs) from distal epithelial progenitor cells after alveolar injury is vital for restoring the epithelial barrier function^2,3^. Following injury, AECIIs act as facultative stem cells of the alveolus and regenerate the gas exchange surface through proliferation and differentiation into AECIs^2^. This progenitor function is partly mediated by Wnt-responsive, Axin2⁺ AECIIs^4^ through injury-associated Krt8⁺ transitional states, shaped by inflammatory cues such as interleukin-1β^5–8^. In addition, bronchioalveolar stem cell (BASCs), located at the bronchioalveolar duct junction, contribute to lung epithelium regeneration exhibiting dual airway and alveolar differentiation potential, depending on the type and severity of the injury^3,9,10^. Notably, in response to bleomycin-induced injury and severe influenza infection, BASCs differentiate into AECIIs and contribute to alveolar epithelial regeneration^3,9^.

AECIIs represent the principal source of granulocyte-macrophage colony-stimulating factor (GM-CSF) in the lung that is required for the differentiation and maintenance of alveolar macrophages^11^. GM-CSF signals via a heterodimeric receptor composed of the ligand-specific α-chain and the common β-chain (GM-CSFRβ), with downstream signaling in alveolar macrophages being essential for terminal differentiation, surfactant clearance, and immune regulation^12,13^. Consistently, disruption of this receptor axis impairs alveolar macrophage functions and causes hereditary pulmonary alveolar proteinosis, underscoring its essential role in maintaining alveolar homeostasis^14,15^. Due to its established role as a myeloid growth factor, and in shaping the alveolar macrophage compartment, GM-CSF has been suggested to impact epithelial maintenance and repair. Early studies demonstrated that epithelial GM-CSF may promote alveolar epithelial cell proliferation and impact lung architecture, suggesting additional functions beyond immune cell modulation^16,17,18^. Reciprocal bone marrow transplantation experiments using WT and *Csf2^⁻/⁻^* (global GM-CSF knockout) mice demonstrated that GM-CSF derived from lung epithelial cells, rather than from immune cells, is critical for surviving lethal IAV infection^19^. However, the direct role of GM-CSF in the activation of epithelial stem cell niches following IAV infection remained unclear.

In the injured lung, activation of AMP-activated protein kinase (AMPK) has been associated with a metabolic shift toward catabolic processes and can limit epithelial cell proliferation^20^. AMPK is a central energy sensor activated under conditions of metabolic stress, such as hypoxia, inflammation, and nutrient deprivation and acts to restrict anabolic growth and cell cycle progression^21,22^.

In this study, we identify GM-CSF as a niche-derived signal selectively produced by distal epithelial progenitor populations upon lung injury. Using *in vivo* injury models, *ex vivo* lung organoids, and primary human epithelial cells, we show that GM-CSF promotes progenitor cell survival, proliferation, and alveolar regeneration. Mechanistically, GM-CSF inhibits AMPK activation, licensing mTORC1 signaling in a GM-CSF receptor-dependent manner, linking cytokine signaling to metabolic control of epithelial progenitor cell function. In the context of IAV-induced lung injury, where epithelial metabolism is reprogrammed through AMPK activation, these findings position GM-CSF as a key regulator of epithelial regeneration and suggest its potential as a therapeutic target during acute injury.

## Results

### GM-CSF expression increases after IAV infection in epithelial progenitor cells and is associated with epithelial cell proliferation and barrier repair

To uncover the epithelial cell population(s) responsible for GM-CSF expression and release during IAV infection, AECs, BASCs and airway epithelial cells were isolated from the lungs of mock- and IAV-infected mice **(Sup. Figure 1A)**. While GM-CSF was not detected in airway epithelial cells, AECs and BASCs showed elevated mRNA levels at 4 and 7 days post-infection (dpi) **(Figure 1A)**. Next, GM-CSF expression in AECIs and AECIIs was evaluated separately. Only AECIIs upregulated GM-CSF implying that epithelial GM-CSF expression is confined to distal epithelial progenitor cell compartments upon IAV infection **(Figure 1B and Sup. Figure 1B)**. No significant expression was detected in lung mesenchymal cells (EpCAM^neg^CD45^neg^Sca-1^+^PDGFRa^+^). To investigate GM-CSF effects on epithelial progenitor cell-driven lung repair in WT, *Csf2^-/-^*, and SPC-GM (GM-CSF over-expressing) mice, we first conducted experiments to adjust the IAV infection doses within these groups to account for their different susceptibilities to IAV infection driven by lack of alveolar macrophages and cDC1 in C*sf2^-/-^* mice and by increased lung myeloid host defense in SPG-GM mice, compared to WT, respectively^13,19^. Comparable viral titers were achieved at 3 dpi, the time point at which IAV titers typically peak^23^, when WT, *Csf2^-/-^*, and SPC-GM mice were infected with 500, 100, and 2000 foci forming units (FFU), respectively **(Figure 1C)**. At 7 dpi, alveolar leakage values as a marker for alveolar barrier function were also equivalent among the mouse strains, further demonstrating that similar levels of lung injury were reached across genotypes allowing for comparative analyses of epithelial repair in the course after 7 dpi **(Figure 1D)**. Using these standardized conditions, distal epithelial progenitor cell proliferation and apoptosis were evaluated. Following IAV infection, the amount of AECIIs as well as AECII and BASC proliferation were significantly elevated in WT and SPC-GM mice when compared to *Csf2^-/-^* mice **(Figure 1E and Sup. Figure 1C-D)**. Conversely, the proportion of apoptotic AECIIs and BASCs was increased in *Csf2^-/-^* mice when compared to infected WT and SPC-GM mice **(Figure 1F)**. Although similar levels of infection and injury were observed at 3 and 7 dpi **(Figure 1C and D)**, elevated alveolar leakage levels persisted in *Csf2^-/-^* mice at 10 dpi **(Figure 1G)**. Moreover, structural alterations of the lung epithelium were similar in WT and *Csf2^-/-^* mice at d7 but failed to decline in *Csf2^-/-^*mice at 14 dpi, as evidenced by increased mean linear intercept (MLI) values, indicative of enlarged alveolar airspaces and impaired restoration of alveolar architecture **(Figure 1H-I and Sup. Figure 1E)**. These findings suggest that sufficient GM-CSF expression in distal epithelial progenitor cell niches in response to viral injury is needed for tissue repair by inducing their proliferation and survival in an autocrine manner.

**Figure 1.**
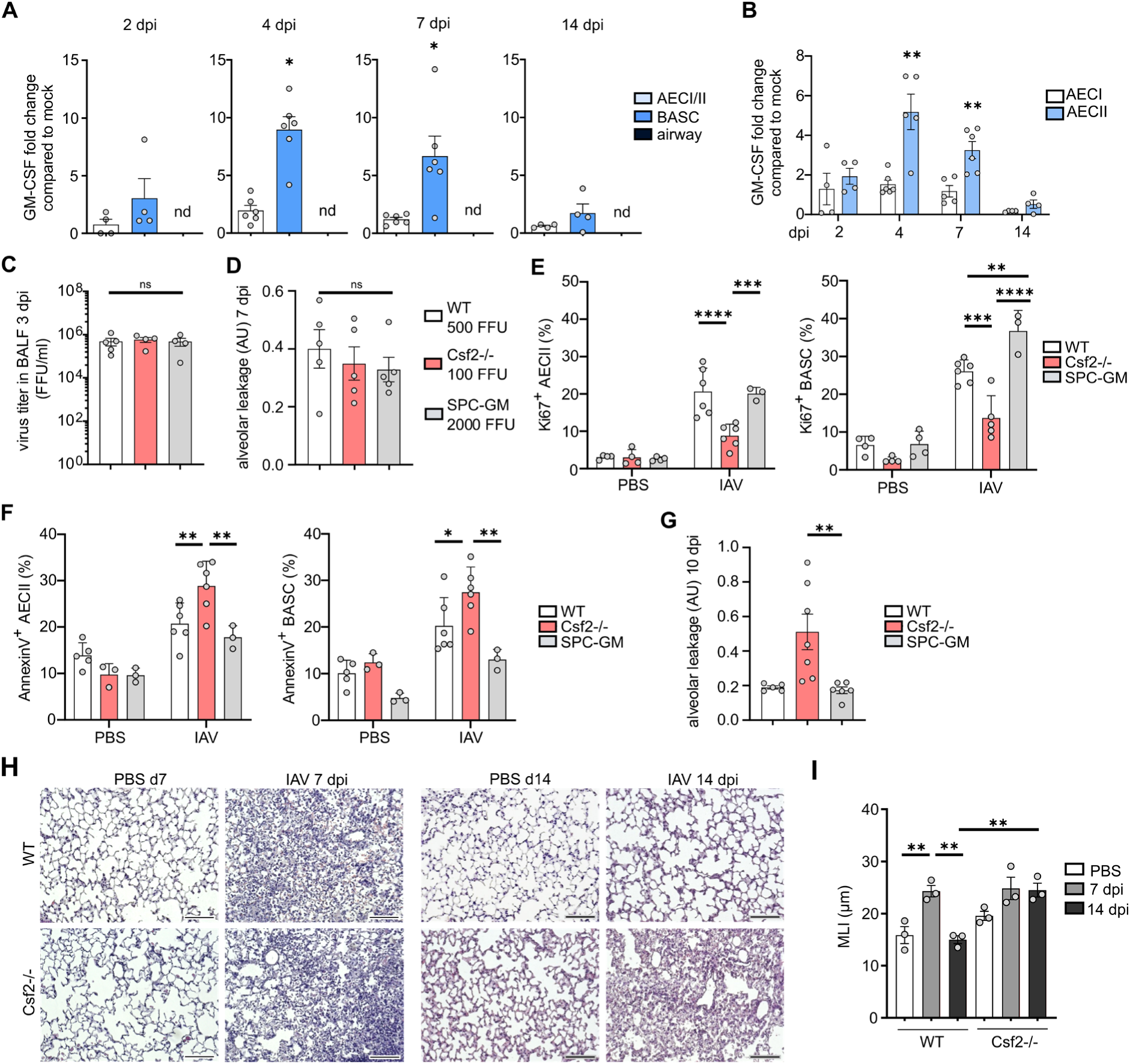
GM-CSF expression is upregulated in epithelial progenitor cells of the distal lung and is associated with epithelial cell proliferation and lung repair. **(A-B)** GM-CSF gene expression at 2, 4, 7, and 14 dpi in **(A)** AECs, BASCs, and airway epithelial cells (n=4-6, independent two-tailed Student’s t-test) and **(B)** in AECI and AECII subpopulations (n=4-5, independent two-tailed Student’s t-test). **(C)** Virus titers in bronchoalveolar lavage fluid (BALF) isolated from WT (500 FFU), *Csf2^-/-^* (100 FFU), and SPC-GM (2000 FFU) mice at 3 dpi (n = 4-5, one-way ANOVA followed by Tukey’s post hoc test). **(D)** Alveolar albumin leakage in WT, *Csf2^-/-^*, and SPC-GM mice at 7 dpi (adjusted doses, n=5, one-way ANOVA followed by Tukey’s post hoc test). **(E)** Ki67^+^ proliferating AECIIs and BASCs in WT, *Csf2^-/-^*, and SPC-GM mice at 7 dpi analyzed by FACS (adjusted doses, n=3-6, one-way ANOVA followed by Sidak’s post hoc test). **(F)** Annexin^+^ apoptotic AECIIs and BASCs in WT, *Csf2^-/-^*, and SPC-GM mice at 7 dpi analyzed by FACS (adjusted doses, n=3-6, one-way ANOVA followed by Sidak’s post hoc test). **(G)** Alveolar albumin leakage in WT, *Csf2^-/-^*, and SPC-GM mice at 10 dpi (adjusted doses, n=5-7, one-way ANOVA followed by Tukey’s post hoc test). **(H)** Transmission microscopy of lung sections (H&E) from WT (500 FFU) and *Csf2^-/-^* (100 FFU) mice at 7 and 14 dpi. Scale bar: 150 µm. **(I)** Analysis of MLI (µm) of eCadherin-stained lung sections from WT (500 FFU) and *Csf2^-/-^* (100 FFU) mice at 7 and 14 dpi versus untreated PBS controls (n=3, two-way ANOVA followed by Sidak’s post hoc test), Graphs show means ± SEM; *=p<0.05; **=p<0.005; ***=p<0.001; ****=p<0.0001.

### GM-CSF signaling is crucial for BASC proliferation, AECII expansion, and alveolarization in bronchoalveolar lung organoids

To validate that GM-CSF signaling contributes to BASC proliferation and subsequent formation of alveoli with AECIIs and AECIs *ex vivo*, we employed an established BASC-derived bronchoalveolar lung organoid (BALO) model, exhibiting proximal-to-distal epithelial differentiation within airways and alveoli **(Figure 2A)**^24^. First, BASC proliferation and organoid formation in response to GM-CSF signaling was investigated in BALO cultures from WT and *Csf2^-/-^* BASCs from non-infected mice. BALO cultures lacking GM-CSF failed to grow and did not develop differentiated BALO structures, indicating that GM-CSF signaling is required for organoid development **(Figure 2B-D).** To further assess the role of GM-CSF/GM-CSF receptor signaling in BASC differentiation into AECIIs and subsequent alveoli formation, BASCs from non-infected WT and *Csf2rb^-/-^* mice were cultured for 21 days. WT BASCs formed BALOs containing multiple alveoli composed of AECIIs and AECIs **(Figure 2E)**. In contrast, *Csf2rb^-/-^* BASCs generated organoids with significantly fewer alveoli when compared to WT controls, implying impaired AECII proliferation and differentiation into AECIs **(Figure 2F)**. Endogenous GM-CSF release was quantified in the supernatants of BALO,cMy and GM-CSF was detectable in the supernatants of WT and *Csf2rb^-/-^* but not in *Csf2^-/-^* cultures **(Figure 2G)**. Since the GM-CSFRβ also forms part of the IL-3 and IL-5 receptor complexes and is required for their signal transduction^25^, these cytokines were additionally measured. Neither cytokine was detectable, confirming that the observed effects in *Csf2rb^-/-^* BALOs are attributable only to GM-CSF signaling **(Sup. Figure 2A and B)**. To further assess whether BALO development, particularly alveolarization, could be rescued in BALO lacking endogenous GM-CSF, *Csf2^-/-^*culture media were supplemented with recombinant GM-CSF **(Sup. Figure 2C)**. Addition of GM-CSF not only enhanced organoid growth but also promoted formation of alveoli, confirming that AECII proliferation in BALO is dependent on GM-CSF signaling **(Figure 2H-K)**. These findings indicate that GM-CSF/GM-CSFRβ signaling drives BASC proliferation and alveolarization with emergence of AECIIs in BALO *ex vivo*.

**Figure 2.**
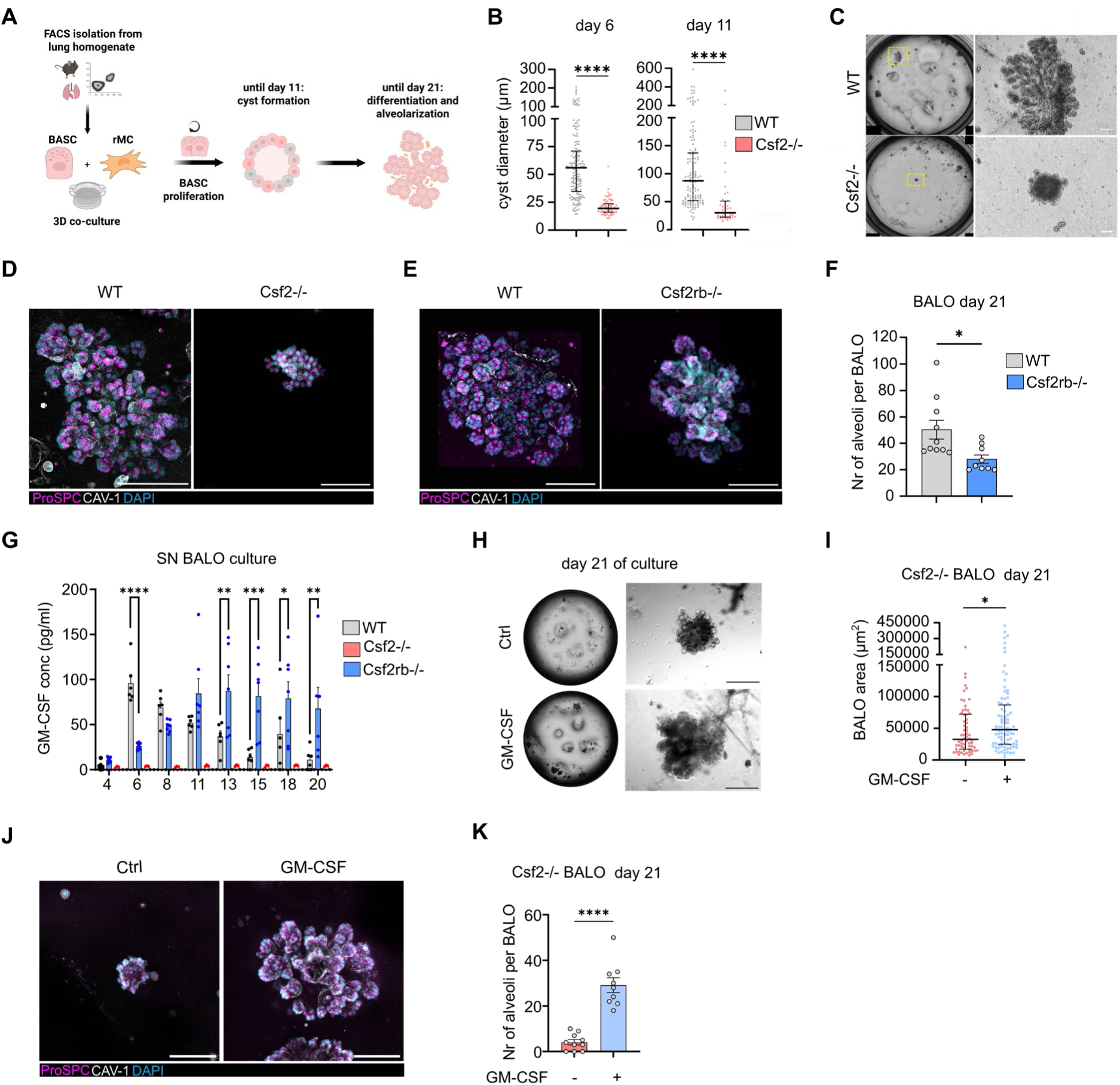
GM-CSF signaling is crucial for BASC proliferation and BALO formation *ex vivo*. **(A)** Scheme of BALO development. **(B)** Cyst diameter at day 6 and 11 of culture in WT and *Csf2^-/-^* BALO cultures (n=3 biological replicates, median with interquartile ranges is shown, Mann-Whitney test). **(C)** Representative transmission light and **(D)** confocal microscopy images of WT and *Csf2^-/-^* BALOs stained for ProSPC (AECIIs), CAV-1 (AECIs), and DAPI at day 21 of culture. Scale bar: 100 µm. **(E)** Confocal microscopy images and **(F)** analysis of the number of alveoli of WT and C*sf2rb^-/-^* BALOs stained for ProSPC (AECIIs), CAV-1 (AECIs), and DAPI at day 21 of culture. Scale bar: 100 µm (day 21, n=3 technical replicates, 3-4 BALOs per well were analyzed, independent two-tailed Student’s t-test). **(G)** GM-CSF concentration in supernatant of WT, *Csf2-/-*, and *Csf2rb^-/-^* BALOs by ELISA (n=3-7 biological replicates, 2-way ANOVA followed by Tukey’s post hoc test). **(H)** Transmission microscopy images of *Csf2^-/-^* BALOs treated with recombinant GM-CSF (50-100 pg/ml) vs. PBS. Scale bar: 500 µm. **(I)** (Bronchio)alveolar lung organoid size (area) in *Csf2^-/-^* cultures treated with GM-CSF vs. PBS (n=3 technical replicates, median with interquartile ranges are shown, Mann-Whitney test). **(J)** Representative confocal images of *Csf2^-/-^* BALOs treated with GM-CSF vs. PBS stained for ProSPC (AECIIs), CAV-1 (AECIs), and DAPI. Scale bar: 100 µm. **(K)** Alveoli count per *Csf2^-/-^* BALO (day 21, n=3 technical replicates, 3-4 BALOs per well were analyzed, independent two-tailed Student’s t-test). Graphs show means ± SEM unless otherwise indicated. *=p<0.05; **=p<0.005; ***=p<0.001; ****=p<0.0001.

### GM-CSF-driven distal epithelial progenitor cell proliferation is mediated by inhibition of AMPK signaling

To unravel the underlying molecular mechanisms by which GM-CSF drives distal epithelial progenitor cell proliferation upon viral injury, RNA-Sequencing of BASCs sorted from the lungs of IAV-infected and mock-treated WT and *Csf2^-/-^* mice was performed **(Sup. Figure 3A)**. Genes related to metabolic pathways such as glycolysis and oxidative phosphorylation were differentially expressed in BASCs from WT IAV-infected versus *Csf2^-/-^* IAV-infected mice **(Figure 3A)**. Our previous studies showed that AMPK (5′-adenosine monophosphate-activated protein kinase), a master regulator of cellular metabolism, is upregulated in AECIIs during IAV infection^26,27^. We hypothesized that GM-CSF may counteract AMPK-dependent metabolic reprogramming. To test this, AECIIs isolated from WT, *Csf2^⁻/⁻^,* and *Csf2rb^⁻/⁻^* mice were infected with IAV *ex vivo* in the presence or absence of a high dose of recombinant GM-CSF to ensure maximal receptor stimulation. IAV infection induced AMPK activation, whereas exogenous GM-CSF suppressed this response in WT and *Csf2^⁻/⁻^* AECIIs, but not in GM-CSF receptor-deficient *Csf2rb^⁻/⁻^* AECIIs **(Figure 3B-D)**. AMPK activation stalls cell proliferation by negatively regulating mTORC1 signaling^26,28^. As an indicator of mTORC1 activity, we quantified phosphorylation of ribosomal protein S6 kinase (phospho-p70S6K), a key downstream target and effector of mTORC1^29^. Addition of recombinant GM-CSF to IAV-infected AECIIs enhanced mTORC1 activity in WT and *Csf2^-/-^* but not *Csf2rb^-/-^* AECII, suggesting that GM-CSF-mediated AMPK inhibition unleashes mTORC1 signaling in a GM-CSFRβ-dependent manner **(Figure 3E-G and Sup. Figure 3B)**. To confirm GM-CSF-mediated effects on AMPK-mTORC1 signaling during viral infection *in vivo*, AECIIs were isolated from mock- and IAV-infected mice 4 days after infection, a time point when epithelial GM-CSF expression peaks **(Figure 1A-B)**. In line with our hypothesis, while AMPK signaling was significantly upregulated in AECIIs from IAV-infected *Csf2^-/-^* and *Csf2rb^-/-^*mice when compared to WT, mTORC1 activity was inhibited **(Figure 3H-I)**. mTORC1 activation promotes cell proliferation by upregulating cell cycle-related genes, including *c-Myc* and cyclin D1 (*Ccnd1*).^30^ Accordingly, WT AECIIs and BASCs isolated from IAV-infected mice at 7 dpi exhibited *c-Myc* upregulation whereas their *Csf2^-/-^* counterparts did not **(Figure 3J)**. Treatment of isolated *Csf2rb^-/-^*AECIIs with exogenous GM-CSF failed to promote *Ccnd1* and *c-Myc* expression **(Figure 3K and Sup. Figure 3C)**. Accordingly, BASCs isolated from IAV-infected *Csf2rb^-/-^* mice at 7dpi showed significantly lower organoid forming efficiency (OFE) compared to WT **(Figure 3L and Sup. Figure 3D-E)**. To assess relevance of these findings in a human model, primary isolated human AECIIs were infected with IAV in the presence or absence of recombinant GM-CSF. AMPK activation after IAV infection was inhibited by addition of exogenous GM-CSF **(Figure 3M)**. Moreover, GM-CSF addition led to mTORC1 signaling activation in IAV-infected human AECIIs **(Figure 3N)**. These results propose that, in response to IAV infection, epithelial GM-CSF antagonizes AMPK signaling, thereby licensing mTORC1 activation and promoting proliferation of distal epithelial progenitor cells.

**Figure 3.**
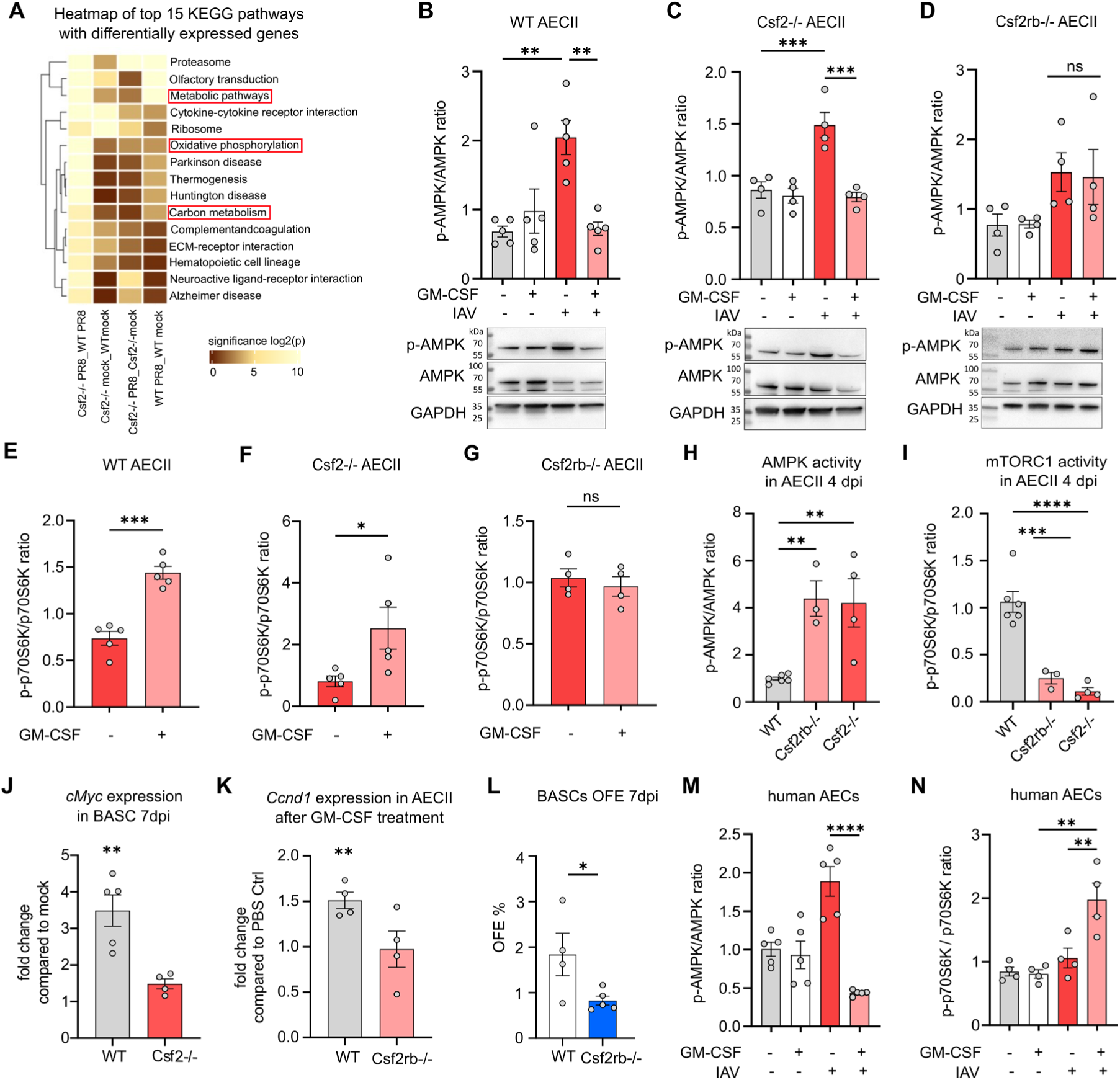
GM-CSF-mediated distal epithelial progenitor cell proliferation after IAV infection is driven by regulation of AMPK signaling. **(A)** Heatmap of KEGG pathways with differentially expressed genes in BASCs obtained from WT and *Csf2^-/-^* IAV-infected and PBS mice at 7 dpi (n=4-5). -Log(p) values range between 0 (black) and 10 (yellow) with high values indicating higher significance. **(B-D)** AMPK activity (phospho-AMPK/AMPK) in primary murine AECs from **(B)** WT, (C) *Csf2^-/-^*, and (D) *Csf2rb^-/-^* mice at 24 h post *ex vivo* IAV infection (MOI 0.1) and GM-CSF treatment (50 ng/ml) analyzed by Western blot (n=4-5, one-way ANOVA followed by Tukey’s post hoc test). **(E-G)** mTORC1 activity (phospho-p70S6K/p70S6K) in primary murine AECs from **(E)** WT, (F) *Csf2^-/-^*, and (G) *Csf2rb^-/-^* mice 24 h post *ex vivo* IAV infection (MOI 0.1) and GM-CSF treatment (50 ng/ml) analyzed by Western blot (n=4-5, independent two-tailed Student’s t-test). **(H)** AMPK and **(I)** mTORC1 activity in AECIIs from WT, *Csf2^-/-^*, and *Csf2rb^-/-^* mice at 4 dpi analyzed by Western blot (n=3-6, one-way ANOVA followed by Tukey’s post hoc test). (J) *c-Myc* expression in BASCs (7 dpi) from WT and *Csf2^-/-^* mice analyzed by qPCR (n=4-5, independent two-tailed Student’s t-test). (K) *Ccnd1* expression in AECIIs treated with GM-CSF (50 ng/ml) *ex vivo* analyzed by qPCR (n=4, independent two-tailed Student’s t-test). **(L)** OFE (%) of BALOs derived from WT (500 FFU) or *Csf2rb^-/-^* (100 FFU) BASCs of IAV-infected mice at 7 dpi (n=4-5 technical replicates, one-way ANOVA followed by Tukey’s post hoc test)**. (M)** AMPK and **(N)** mTORC1 activity in human AECs *ex vivo* after 45 min (AMPK) and 30 min (mTORC1) IAV infection (MOI 1) and human GM-CSF treatment (100 ng/ml) analyzed by Western blot (AMPK Western blot n=5, mTORC1 Western blot n=4, one-way ANOVA with Tukey’s post hoc test). Graphs show means ± SEM; *=p<0.05; **=p<0.005; ***=p<0.001; ****=p<0.0001.

### GM-CSF-AMPK-mTORC1 signaling drives distal lung epithelial progenitor cell proliferation and alveoli formation *ex vivo*

To validate that GM-CSF-AMPK-mTORC1 signaling is crucial for proliferation of lung progenitor cells, BALO (derived from BASCs) and alveolospheres (generated from AECIIs) were treated with the AMPK modulators AICAR (AMPK activator) and Compound C (AMPK inhibitor) **(Sup. Figure 4A and Sup. Figure 5A)**. OFE of BALOs and alveolospheres were reduced after treatment with AICAR **(Figure 4A, Sup. Figure 4B and 5B-C)**. In contrast, organoid numbers were significantly increased after Compound C treatment **(Figure 4B and Sup. Figure 4C and 5D-E)**. Notably, AICAR-treated BALOs displayed reduced AECII signal and yielded fewer alveoli when compared to PBS and Compound C-treated cultures **(Figure 4C-D)**. Impaired BALO formation upon AICAR treatment was reversed by simultaneous addition of the mTORC1 activator, MHY1485 **(Figure 4E and Sup. Figure 4D)**. Of note, addition of exogenous GM-CSF to WT cultures not only led to AMPK inhibition but also increased AECII signal in BALO **(Figure 4F-G and Sup. Figure 4E)**. Accordingly, addition of Compound C or MHY1485 significantly improved OFE in GM-CSF-deficient BALO **(Figure 4H and Sup. Figure 4F-H)**. In addition, AMPK activation by AICAR reduced mTORC1 activity, whereas subsequent exogenous GM-CSF treatment restored mTORC1 activation and alveoli formation in BALO **(Figure 4I-J)**. This mechanism was confirmed in murine alveolospheres **(Sup. Figure 5A-E)** and in primary human AECII-derived alveolospheres **(Figure 4K-L, Sup. Figure 6A-C)**. These data suggest that GM-CSF inhibits AMPK signaling in lung epithelial progenitor cells, thereby licensing mTORC1 signaling and enhancing their proliferation and subsequent alveolarization in lung organoids.

**Figure 4.**
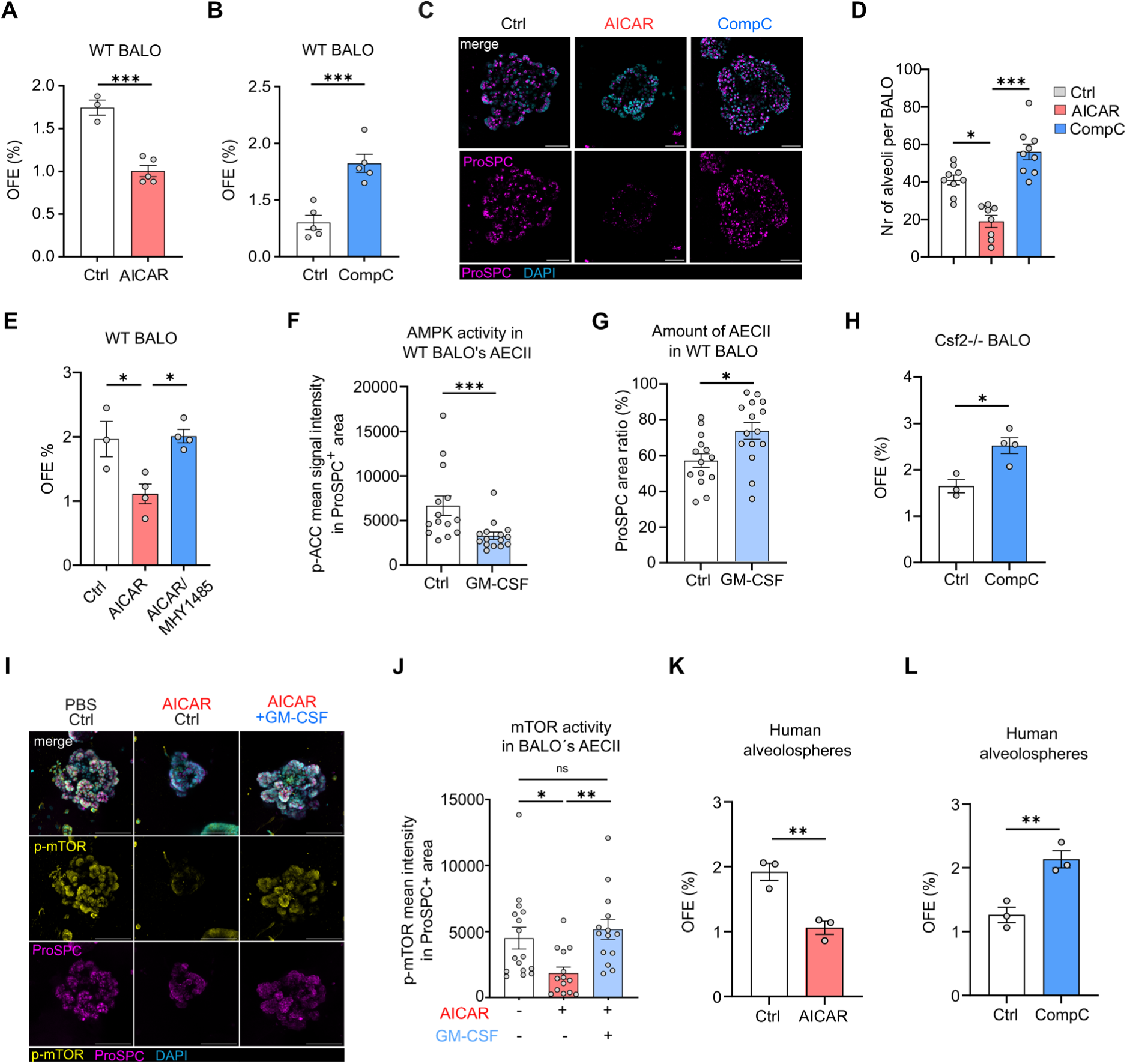
GM-CSF-AMPK-mTORC1 signaling regulates distal epithelial progenitor cells proliferation and lung organoid formation. **(A-B)** OFE (%) of WT BALO cultures treated with **(A)** AICAR vs. PBS (n=3-5) and **(B)** Compound C vs. DMSO (day 10-21) (n=5 technical replicates, independent two-tailed Student’s t-test). **(C)** Representative confocal images of WT BALOs treated with AICAR or Compound C, stained for ProSPC (AECIIs/BASCs) and CAV-1 (AECIs). Scale bar: 50 µm. **(D)** Number of alveoli per BALO treated with AICAR, Compound C vs PBS control (day 21, n=3 technical replicates, 3-4 BALOs per well were analyzed, one-way ANOVA followed by Tukey’s post hoc test). **(E)** OFE (%) of BALOs treated with AICAR or AICAR + MHY1485 vs. PBS/DMSO (day 10-21) (n=3-4 technical replicates, one-way ANOVA followed by Tukey’s post hoc test). **(F)** AMPK activity (phospho-ACC MFI) (Mann-Whitney test) and **(G)** ProSPC^+^ area (%) in BALOs (day 12) treated with GM-CSF (50 ng/ml) vs PBS Ctrl from day 6 to 12 (n=3 technical replicates, 4-5 images per well were analyzed, independent two-tailed Student’s t-test). **(H)** OFE (%) of *Csf2^-/-^* BALOs treated with Compound C vs. DMSO (n=3-4 technical replicates, independent two-tailed Student’s t-test). **(I-J)** BALOs treated with AICAR (0.5 mM) (day 4-8) followed by GM-CSF (100 ng/mL) (day 8-12) vs. PBS. **(I)** Representative confocal images of BALOs stained for ProSPC and p-(S2448)-mTOR (indicating mTORC1 activity). Scale bar: 50 µm. **(J)** mTORC1 activity (p-mTOR MFI) in ProSPC^+^ AECII/BASC in BALOs (n=3 technical replicates, 4-5 BALOs per well were analyzed, Kruskal-Wallis test followed by Dunn’s post hoc test) **(K-L)** OFE (%) of human alveolospheres treated with **(K)** AICAR vs. PBS and **(L)** Compound C vs. DMSO (day 21-28) (n=3 technical replicates, independent two-tailed Student’s t-test). Concentrations used: AICAR (0.5 mM), Comp C (0.5-1 µM) and MHY1485 (1 µM). Graphs show means ± SEM; *=p<0.05; **=p<0.005; ***=p<0.001; ****=p<0.0001.

### GM-CSF-induced distal lung epithelial progenitor cell proliferation after viral injury depends on AMPK signaling *in vivo*

To validate our findings *in vivo*, GM-CSF effects on AMPK signaling and lung barrier restoration were examined in the IAV-induced lung injury model. Infected WT mice were treated with either AICAR, Compound C or PBS at 3 and 5 dpi via intrapulmonary deposition **(Figure 5A)**. At 7 dpi, AMPK activation resulted in increased AECII and BASC apoptosis compared to Compound C or control treatment, and Compound C treatment reduced BASC apoptosis compared to untreated controls **(Figure 5B-C Sup. Figure 7)**. Moreover, AECIIs and BASCs from AICAR-treated mice showed reduced proliferative capacity compared with Compound C- or control-treated mice, and Compound C increased the number of ProSPC+ cells **(Figure 5D-H and Sup. Figure 8)**. Consistent with these findings, AICAR-treated mice maintained elevated alveolar albumin leakage up to 10 dpi **(Figure 5I)**.To corroborate these findings in a GM-CSF-deficient scenario, *Csf2^-/-^* mice were infected with IAV and treated with Compound C or PBS to assess if AMPK inhibition can restore epithelial progenitor cell function and promote lung repair **(Figure 5J)**. After 7 dpi, mice treated with Compound C significantly increased the number of proliferative AECIIs and BASCs while reducing apoptosis in both distal epithelial progenitor populations **(Figure 5K-N)**. Alongside, Compound C treatment reduced epithelial barrier dysfunction **(Figure 5O)**. These data suggest that pharmacological inhibition of AMPK following virus-induced injury promotes the survival and proliferation of distal epithelial progenitor cells, enhancing lung barrier restoration. These effects are particularly evident in GM-CSF-deficient mice, where impaired repair responses are more severe but can be significantly improved by Compound C-mediated AMPK signaling inhibition.

**Figure 5.**
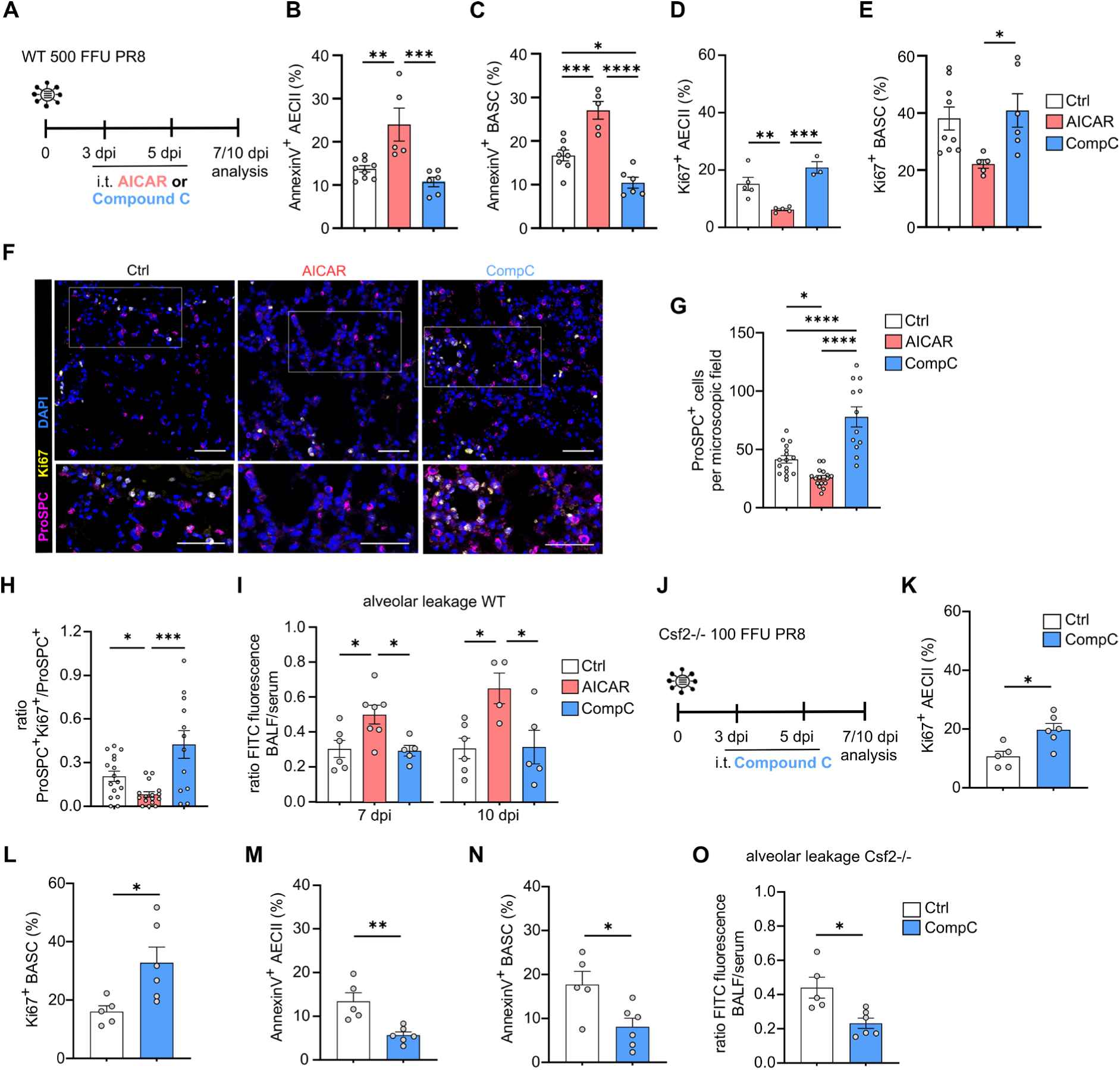
GM-CSF-driven distal lung epithelial progenitor cell proliferation after viral injury is dependent on AMPK signaling *in vivo*. **(A)** Experimental setup: C57BL/6 WT mice (500 FFU PR8) were treated intratracheally with AMPK activator (AICAR; 500 mg/kg bodyweight) or AMPK inhibitor (Compound C; 20 mg/kg bodyweight) vs. control at 3 and 5 dpi. Analyses were performed at day 7 and 10 pi. **(B-C)** Apoptotic (AnnexinV^+^) **(B)** AECIIs (n=5-9) and **(C)** BASCs (n=5-8) at 7 dpi in WT analyzed by FACS (one-way ANOVA followed by Tukey’s post hoc test). **(D-E)** Proliferating (Ki67^+^) **(D)** AECIIs (n=3-5) and **(E)** BASCs (n=5-9) at 7 dpi in WT by FACS (one-way ANOVA followed by Tukey’s post hoc test). **(F)** Confocal microscopy images of lung sections from WT 7 dpi stained for ProSPC, Ki67, and DAPI. Scale bar: 50 µm (upper panel) and 20 µm (lower panel). **(G)** ProSPC^+^ cell count per microscopic field and **(H)** Ki67^+^ProSPC^+^ / ProSPC^+^ ratio were determined (n=3-4 mice, 4 images per mouse were analyzed, one-way ANOVA followed by Tukey’s post hoc test). **(I)** Alveolar leakage (ratio FITC albumin in BALF / serum) at 7 dpi (n=5-7) and 10 dpi (n=4-6, one-way ANOVA followed by Tukey’s post hoc test). **(J)** Experimental setup: *Csf2^-/-^* mice (100 FFU PR8) were treated intratracheally with AMPK inhibitor Compound C vs. control at 3 and 5 dpi. **(K-L)** Amount of proliferating (Ki67^+^) **(K)** AECIIs and **(L)** BASCs at 7 dpi analyzed by FACS (n=5-6, independent two-tailed Student’s t-test). **(M-N)** Amount of apoptotic (AnnexinV^+^) **(M)** AECIIs and **(N)** BASCs at 7 dpi by FACS (n=5-6, independent two-tailed Student’s t-test). **(O)** Alveolar leakage at 10 dpi (n=5-6, independent two-tailed Student’s t-test). Graphs show means ± SEM; *=p<0.05; **=p<0.005; ***=p<0.001; ****=p<0.0001.

### Intrapulmonary GM-CSF deposition enhances alveolar epithelial barrier repair after IAV-induced injury

To investigate if GM-CSF treatment would promote distal progenitor cell proliferation and repair, recombinant GM-CSF was applied into the lungs of IAV-infected mice at 3 and 5 dpi. These timepoints were chosen to benefit from both, its anti-apoptotic effect that might prevent alveolar epithelial cell injury, and the proliferative effect on lung progenitor cells aiding in repair following injury **(Figure 6A)**. Epithelial barrier dysfunction was significantly decreased after GM-CSF treatment, resulting in higher oxygen saturation levels at 10 dpi when compared to PBS controls **(Figure 6B-C)**. Concomitantly, AECII numbers were increased in GM-CSF-treated mice, accompanied by a reduction in the proportion of damaged lung areas **(Figure 6D-G).** Accordingly, lung tissue architecture was improved in treated mice as evidenced by decreased MLI values, indicative of enhanced (neo-) alveolarization **(Sup. Figure 9A-B)**. The improved epithelial barrier function and increased oxygenation following GM-CSF treatment suggested that expanded AECIIs transdifferentiated into AECIs to re-establish the gas exchange surface. SPC^CreERT2;tdTomato^ mice were infected with IAV and treated with GM-CSF or control at 3 and 5 dpi allowing AECII lineage tracing by quantification of tdTomato^+^ AECIs **(Figure 6H, Sup. Figure 9C)**. In fact, the proportion of tdTomato^+^ of total AECI was increased in response to GM-CSF treatment **(Figure 6I)**. Moreover, GM-CSF treatment increased the abundance of tdTomato⁺RAGE⁺ AECIs in precision-cut lung slices, with the most pronounced effect observed within the defined repair-associated regions (Zone 2), whereas no changes were detected within severely damaged regions widely lacking progenitor cells (Zone 1) or non-injured lung regions (Zone 3)^4^ at 14 dpi **(Figure 6J-K, Sup. Figure 9D)**. These results confirm that GM-CSF-driven progenitor cell expansion expansion promotes alveolar epithelial barrier repair via re-establishment of the AECI pool. Furthermore, these findings highlight intrapulmonary GM-CSF delivery as a putative therapeutic approach for hypoxemic acute lung injury in the context of viral infection.

**Figure 6.**
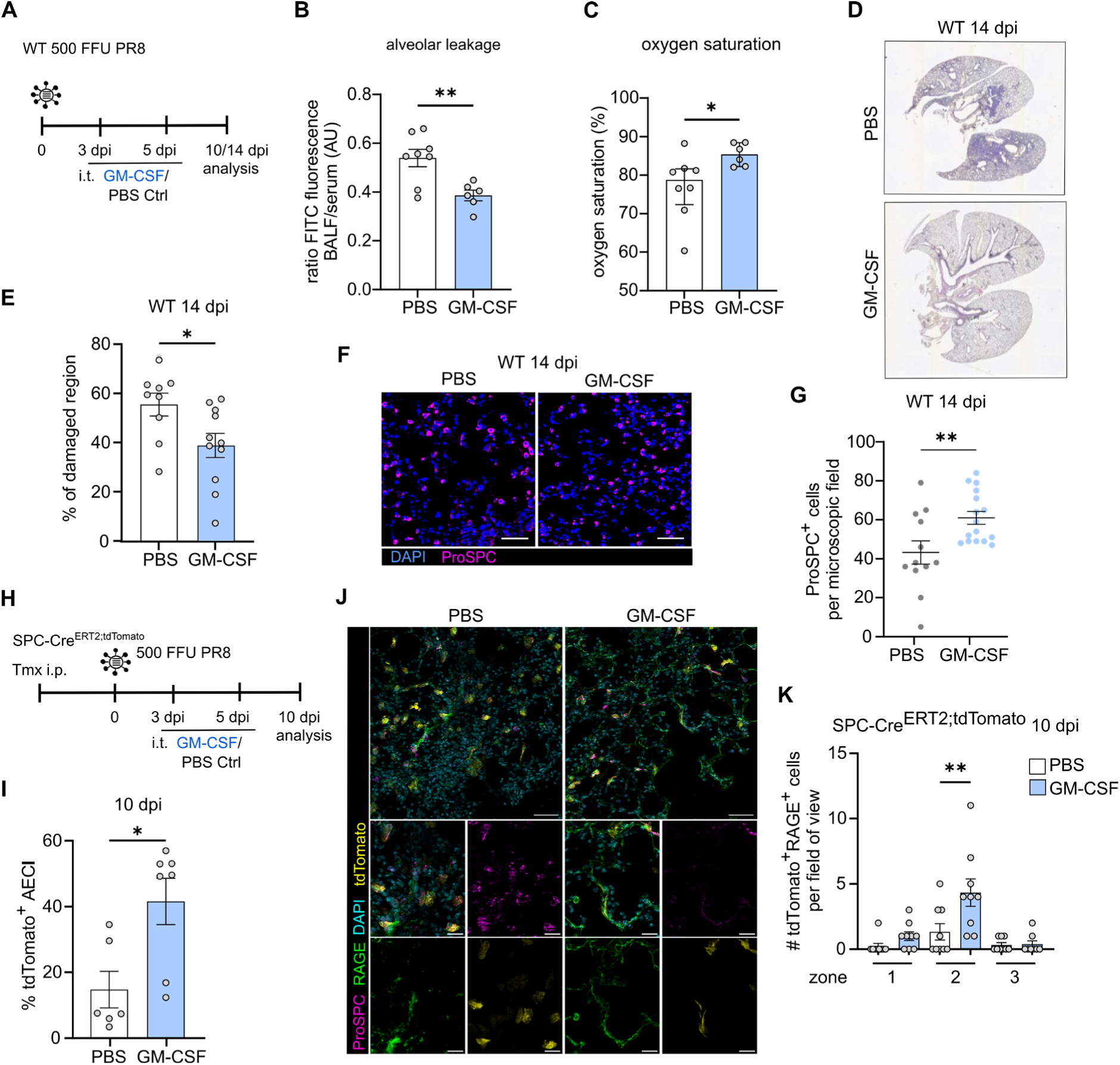
Intrapulmonary GM-CSF administration enhances alveolar epithelial barrier repair after IAV-induced injury. **(A)** C57BL/6 mice were infected with IAV and intratracheally treated with 10 µg GM-CSF versus PBS Ctrl at 3 and 5 dpi. Analyses were performed at 10 and 14 dpi. **(B)** Alveolar leakage (ratio FITC fluorescence BALF/serum) and **(C)** oxygen saturation at 10 dpi (n=6-8, independent two-tailed Student’s t-test). **(D)** Transmission microscopy images of H&E-stained lung sections (14 dpi) and **(E)** analysis of the percentage of damaged regions (n=3-4, 3 slices per mouse lung were analyzed, independent two-tailed Student’s t-test). **(F)** Confocal microscopy images of lung sections (14 dpi) stained for ProSPC and DAPI and **(G)** analysis of ProSPC^+^ cells (AECII and BASC) per microscopic field (n=3-4, 4 images per lung were analyzed, independent two-tailed Student’s t-test). **(H)** Experimental setup: Tamoxifen-induced labeling of AECIIs in SPC-Cre^ERT2^;tdTomato mice, followed by IAV infection (250 FFU, day 0) and i.t. treatment with 10 µg GM-CSF vs. PBS (3 and 5 dpi). Analysis was performed at 10 dpi. **(I)** Amount of tdTomato^+^ AECIs (%) were measured by FACS. **(J)** Vibratome sections were stained for ProSPC, RAGE, and DAPI (scale bar upper panel 50 µm, lower panel 20 µm) **(K)** and the amount of tdTomato+RAGE+ AECIs per field of view was determined (n=3, three images per mouse were analyzed, one-way ANOVA followed by Tukey’s post hoc test). Graphs show means ± SEM; *=p<0.05; **=p<0.005.

## Discussion

We identified that GM-CSF expression in the injured lung is restricted to distal epithelial compartments with progenitor potential, including AECIIs and BASCs. While GM-CSF has been extensively characterized as a cytokine that guides alveolar macrophage identity and function^12,13^, our data extend this paradigm by demonstrating that GM-CSF also functions as a local epithelial niche factor regulating distal progenitor epithelial cell behavior. This dual role positions GM-CSF at the interface between immune regulation and epithelial regeneration, highlighting intrapulmonary delivery as a new therapeutic avenue in IAV-induced ARDS and beyond^31,32^.

A limitation of our *in vivo* models is that GM-CSF effects on immune cells cannot be fully excluded, given its well-established role in macrophage and dendritic cell activation^33^. However, the use of alveolospheres and BALOs lacking myeloid cells demonstrates that GM-CSF directly promotes epithelial proliferation and alveoli formation. These findings clearly establish an epithelial-intrinsic function of GM-CSF independent of its immune-modulatory effects. Importantly, our findings place GM-CSF within a broader, AECII-centered, regenerative program that restores alveolar structure following injury. AECIIs regenerate AECIs through transitional states such as DATPs, which are regulated by inflammatory signals including IL-1β^5^. While these signals are required to initiate repair, persistent inflammation can impair lung epithelial differentiation and lead to defective regeneration^10,34,35^. Our data suggest that GM-CSF promotes expansion of distal epithelial progenitor cells and increases the AECI pool, thereby enhancing alveolar repair. Whether GM-CSF directly regulates the AECII-to-AECI differentiation trajectory or primarily expand the progenitor pool remains unresolved. It is conceivable that GM-CSF acts upstream of transitional states by increasing the pool of competent progenitors, while additional signals fine-tune lineage progression^36^. Further studies are necessary to define how GM-CSF interacts with established regulators of transitional states formation during injury.

A key mechanistic insight of this study is that GM-CSF links cytokine signaling to metabolic programs in epithelial progenitor cells. We demonstrate that GM-CSF inhibits AMPK activation and enables mTORC1 signaling, thereby allowing cells to transition from a stress-adapted state to a proliferative state. AMPK activation is a conserved response to metabolic stress and limits cell growth, whereas mTORC1 promotes anabolic metabolism and proliferation^21,22^. Inflammatory cues and injury-induced stress responses, including those triggered by viral infection, further enhance AMPK signaling in AECs, thereby imposing a metabolic checkpoint that must be overcome for effective regeneration^27^. Stem and progenitor cell fate transitions are increasingly recognized to require metabolic plasticity, with glycolytic and mitochondrial programs being dynamically engaged to support proliferation, lineage specification and differentiation^37,38^. Consistent with this, AMPK signaling disturbances have been linked to impaired alveolar repair and defective epithelial regeneration^36^. Thus, GM-CSF may act as a temporal regulator of epithelial repair by promoting an anabolic, mTORC1-permissive state during progenitor expansion, while subsequent differentiation may require additional metabolic remodeling, including restoration of mitochondrial oxidative programs.

The molecular connection between GM-CSF receptor activation and AMPK inhibition, however, remains to be defined. GM-CSF receptor signaling activates the PI3K-AKT pathway via the common β-chain, which can promote mTORC1 activation and oppose catabolic signaling^39,40^. It is therefore plausible that GM-CSF counteracts the known AMPK effects also indirectly through PI3K-AKT-mTORC1 signaling. Additionally, GM-CSF has been shown to impact mitochondrial metabolism, suggesting that it may reduce metabolic stress signals that trigger AMPK activation^41^. These findings highlight a previously unrecognized role for GM-CSF in coordinating metabolic reprogramming during epithelial repair. In addition to its effects on proliferation, GM-CSF exerts a pronounced anti-apoptotic effect on epithelial progenitor cells^17,18^. This is of particular importance in the context of lung injury, where excessive epithelial cell death can limit regenerative capacity. By preserving AECIIs and BASCs, GM-CSF ensures maintenance of the progenitor pool required for effective repair. This is consistent with previous studies demonstrating survival-promoting effects of GM-CSF through PI3K- and MAPK-dependent pathways^41^.

GM-CSF has been investigated as a therapeutic agent in pulmonary infections due to its ability to enhance host defense through activation of myeloid cells, and to restore or maintain the alveolar macrophage pool ^33^. Clinical applications and RCTs of inhaled GM-CSF in patients with pneumonia and ARDS have demonstrated its potential to improve alveolar macrophage immune function and clinical outcomes^31,32,42^. Our findings extend this framework by demonstrating that GM-CSF directly promotes epithelial repair by enhancing progenitor cell proliferation, alveolarization, and barrier restoration. This dual function of enhancing antimicrobial defense and alveolar macrophage functions, while promoting tissue regeneration has important therapeutic implications, representing a strategy to simultaneously improve pathogen clearance and accelerate alveolar epithelial regeneration via the AMPK-mTOR axis.

### Methods

#### Study approval

All animal experiments were performed at Justus-Liebig University Giessen and were approved by the responsible animal ethics committee and by the local authorities by the state of Hesse (Regierungspräsidium Giessen). Human lung tissue samples were obtained from patients who underwent lobectomy after informed written consent at Justus-Liebig-University Giessen. The approval for the use of human lung tissue samples from the biobank (AZ 10/06, amendment to proposal 31/93) was given in the context of the research center SFB 1021 (project C05) by the Ethics Committee of Justus-Liebig University Giessen.

#### Mice

Mice were housed under specific pathogen-free conditions. C57BL/6 (WT) mice were purchased from Charles River Laboratories. GM-CSF receptor β-deficient (*Csf2rb^-/-^*) [B6.129S1-Csf2rbtm1Cgb/J] and GM-CSF-deficient (*Csf2^-/-^*) [B6.129S-Csf2tm1Mlg/J] knockout mouse strains were purchased from Jackson Laboratories. *Csf2^-/-^* mice have a global knockout of GM-CSF whereas *Csf2rβ^-/-^*mice lack the β-chain of the GM-CSF receptor preventing GM-CSF signaling^13,19^. Transgenic SPC-GM mice [B6.129S1-csf2^tm1mIg(SPC-GM-CSF)^] were received from Dr. Jeffrey Whitsett (University of Cincinnati, OH). These mice are characterized by a global GM-CSF knockout but overexpression of GM-CSF in SPC-positive cells, like AECIIs and BASCs. SPC-GM mice were generated from *Csf2*^-/-^ mice by insertion of a chimeric gene consisting of the GM-CSF sequence controlled by the human SPC promoter^13,16^ _SPCCreERT2;tdTomato mice_ [Sftpc^tm1(cre/ERT2,rtTA)Hap^Gt(ROSA)26Sor^tm9(CAG-tdTomato)Hze^/sbel] were provided by Prof. Dr. Elie El Agha. Sftpc^tm1(Cre/ERT2,rtTA)Hap^ (SPC^Cre-ERT2^) mice were described previously^43^, while Gt(ROSA)26Sor^tm9(CAG-tdTomato)Hze^ (tdTomato) mice were obtained from the Jackson Laboratory (stock number 007905, Bar Harbor, ME, USA). The two mouse lines were crossed to generate SPC^CreERT2^;tdTomato mice where Tamoxifen administration induces tdTomato expression specifically in SPC+ cells^44^. Tamoxifen powder (Sigma-Aldrich, T5648, St. Louis, MO, USA) was dissolved in 100 µl corn oil (Sigma-Aldrich, C8267) and administered via intraperitoneal injection three times (day -18, -16, and -14 before infection) at a concentration of 250 µg/g body weight. Mice were housed under specific-pathogen-free (SPF) conditions with ad libitum access to food and water.

#### Cell culture and primary AEC isolation

Commercially available MDCK cells, primary hAECs, and MRC-5 cells were cultured according to the manufacturer’s instructions. Primary murine AEC isolation was based on a protocol previously described^45,46^. AECs were cultured in growth medium consisting of DMEM (1×) supplemented with 10% fetal calf serum (FCS), 1% penicillin/streptomycin (100 U/mL penicillin and 10 mg/mL streptomycin), and 1% L-glutamine (200 mM).

For the isolation of primary human AECs, lung tissue was mechanically dissociated, washed, and filtered through a 100 μm filter. Cells were enzymatically digested with 2.5% dispase II (in 2 mM Ca²⁺, 1.3 mM Mg²⁺) for 180 min at 37°C, sequentially filtered (100, 40, and 20 μm), and isolated by Ficoll density-gradient centrifugation. Leukocytes were depleted using anti-human CD45 magnetic beads (#130-045-801, Miltenyi Biotech, Germany) according to the manufacturer’s instructions. Human AECs were assessed for purity by flow cytometry (90-98% epithelial cells) and viability by trypan blue exclusion (>95%). Cells were seeded at 3-4.5 × 10⁵ cells/cm² and cultured for 5–6 days to 85-90% confluency before treatment. Human AECs were cultured in Ham’s F-12 Nutrient Mixture (1×) supplemented with 10% FCS, 1% penicillin/streptomycin (100 U/mL penicillin and 10 mg/mL streptomycin), 1% L-glutamine (200 mM), and 1% amphotericin B (250 μg/mL).

#### Organoid culture

Murine alveolospheres and BALOs were generated according to the protocols previously described and cultured up to 21 days^24,44^. The generation of human alveolospheres was conducted according to a previously published protocol^34^. Organoid cultures were treated with the following compounds: AICAR (0.5 mM), Compound C (0.5 and 1 µM), MHY1485 (1 µM), and recombinant murine GM-CSF (50 and 100 pg/mL).

#### In vivo infection and treatments

For *in vivo* infections, 100, 500, or 2000 foci forming units (FFU) of PR8 IAV diluted in PBS were intratracheally applied to mouse lungs. The following compounds were intratracheally applied: AICAR (500 µg/g body weight) diluted in NaCl, Compound C dihydrochloride (20 µg/g bodyweight) and recombinant GM-CSF (10 µg per mouse) both diluted in PBS.

#### Alveolar leakage measurement

Alveolar barrier integrity was assessed by intravenous administration of fluorescein isothiocyanate (FITC)-labeled albumin. Mice were anesthetized by intraperitoneal injection of ketamine hydrochloride (100 mg/kg body weight) and xylazine hydrochloride (16 mg/kg bodyweight) both diluted in 0.09% sterile NaCl, and 1 mg FITC-albumin diluted in 100 µL NaCl was injected into the tail vein. After 45 min, mice were euthanized by cervical dislocation, and blood was collected from the vena cava. BALF was collected in three steps (300, 400, and 500 µl PBS/EDTA) and processed to obtain cell-free supernatant. Blood samples were allowed to clot for 2 h at room temperature in the dark before serum isolation. FITC fluorescence in BALF and 1:10 diluted serum was measured at 525 nm (FLX800 plate reader, BioTek). Alveolar permeability was quantified as the ratio of BALF to serum fluorescence intensity (alveolar leakage = fluorescence BALF / fluorescence blood serum).

#### Ex vivo infection and treatments

In *ex vivo* experiments, murine AECs were infected with the influenza virus strain A/Puerto Rico/8/34 (PR8, H1N1) at an MOI of 0.1 and human AECs at an MOI of 1. At 85-95% confluency, primary AECs were washed with PBS+/+ and inoculated with PR8 IAV in 0.2% BSA/PBS+/+. After 45 min of incubation (37°C, 5% CO₂), the inoculum was replaced by infection medium (10% FCS was replaced by 0.2% BSA). Ex vivo IAV-infected AECs were treated with recombinant murine (50 ng/mL) or human (100 ng/mL) GM-CSF.

#### Virus titration

Virus titers in BALF from IAV infected mice were determined by immunohistochemistry-based plaque assay on 85–90% confluent MDCK cells in 96-well plates in triplicates. Cells were incubated with 50 µL virus dilutions (1:10^1^ to 1:10^8^) for 40 min at 37°C and covered with 100 µL Avicell overlay medium (2xMEM, 50% Avicell 2.5%, 1 µg/mL TPCK-Trypsin) for 24 h. After overlay removal, the cells were fixed with 4% paraformaldehyde (PFA), permeabilized (0.3% Triton-x-100 in PBS-/-), and stained with anti-nucleoprotein primary antibody for 1 h, followed by 1 h incubation with the secondary HRP-linked antibody (ab6789, Abcam). KPL TrueBlue^TM^ peroxidase substrate (seracase) was added to visualize the plaques. To calculate the virus titer, plaques were counted at the microscope (BX41 light microscope, Leica).

#### Western Blot

For Western Blot analysis cultured AECs or FACS-isolated cells were lysed in NP-40 cell lysis buffer (20 mM Tris, 150 mM NaCl, 1 mM EDTA, 1 mM EGTA, 0.5% NP40, 1 mM Orthovanadat) containing protease inhibitor (ThermoFisher) and phosphatase inhibitor cocktails (CellSignaling). The protein concentrations were determined utilizing Bradford Assay (BioRad). Separation of proteins was resolved on an SDS-PAGE and transferred onto PVDF-membranes (BioRad). Membranes were blocked and incubated with the following antibodies (Cell Signaling): AMPKα Rabbit Antibody (2532), phospho-AMPKα (Thr172) Rabbit Antibody (2535), p70 S6 Kinase Rabbit Antibody (9202), phospho-p70 S6 Kinase (Thr389) Rabbit Antibody (9205), GAPDH (14C10) Rabbit mAB (2118), and anti-rabbit IgG HRP-linked Antibody (7074). The bands were detected using the ChemiDoc XRS+ imaging system (BioRad).

#### Flow cytometry and cell sorting

Multicolor flow cytometry and cell sorting were performed using an LSR Fortessa and BD FACSAria III cell sorter with DIVA software (BD Biosciences). Cells obtained from perfused and homogenized mouse lungs (with or without depletion of endothelial cells and leukocytes) were freshly stained with fluorochrome-labeled antibodies for 15 min at 4°C in MACS buffer (1mM EDTA, 0.5% FCS in PBS-/-, pH 7.2). This cell surface staining was done for CD31 (clone MEC 13.3, Biolegend), CD45 (clone 30-F11, BD Biosciences), EpCAM (clone G8.8, Biolegend), CD24 (clone M1/69, Biolegend), Sca-1 (clones E13-161.7 and D7, Biolegend), and T1α (clone 8.1.1, Biolegend). The intracellular staining of Ki67 (clone 16A8, Biolegend) was done subsequently, utilizing the Foxp3/Transcription Factor Staining Buffer Set (ThermoFisher). As negative controls, corresponding isotype antibodies were used. Dead cell staining was done with Sytox^TM^ Blue (ThermoFisher). For AnnexinV staining, cells were resuspended in an AnnexinV binding buffer containing Pacific Blue-labeled AnnexinV (1:20 dilution) (ThermoFisher) and incubated for 15-25 min at 4°C. Ki67^+^ and AnnexinV^+^ AECs and BASCs from *in vivo* infected mice are given as a percentage of the EpCAM^low^ (AEC) or EpCAM^high^CD24^low^Sca-1^+^ (BASC) population (gating strategies adapted from Vazquez-Armendariz et al^24^).

#### ELISA

Organoid culture supernatants were analyzed using a commercially available ELISA kit for mouse GM-CSF (R&D systems) according to the manufacturer’s instructions.

#### Quantitative PCR

For RNA isolation the RNeasy kit (QIAGEN) was utilized, and cDNA synthesis was done as previously described. Quantitative PCR (qPCR) was performed with the iTaq SybrGreen Mix (BioRad) and the AB Step one plus Detection System (Applied Bioscience) or the QuantStudio™ 3 Real-Time PCR System (ThermoFisher). *GAPDH* or *mRPS18* served as the normalization control. Data are presented as fold-change (2^-ΔΔCt^). The following primer pairs were used: *GAPDH* (fwd 5’-CCCCCATGTTTGTGATGGGT-3’; rev 5’-TCTTCTGGGTGGCAGTGATG-3’), *mRPS18* (fwd 5’-CCGCCATGTCTCTAGTGATCC-3’; rev 5’-TTGGTGAGGTCGATGTCTGC-3’), *Csf2* (fwd 5’-CCCCCATGTTTGTGATGGGT -3’; rev 5’-TCTTCTGGGTGGCAGTGATG-3’), *Ccnd1* (fwd 5’-CGTGGCCTCTAAGATGAAGG-3’; rev 5’-CTGGCATTTTGGAGAGGAAG-3’), and *c-Myc* (fwd 5’-ACCACCAGCAGCGACTCTGA-3’; rev 5’-TGGCAGGGGTTTGCCTCTTC-3’).

#### RNA sequencing

BASCs were isolated from lung homogenates of WT and *Csf2^-/-^* mice, either IAV-infected or mock-treated, using FACS. RNA was then extracted from the isolated BASCs, and the samples underwent quality control analysis before proceeding to RNA sequencingFor each condition, 4-5 biological replicates were analyzed and included in the Kyoto Encyclopedia of Genes and Genomes (KEGG) pathway analysis.

#### Histology

For immunofluorescence or H&E staining of lung sections, mouse lungs were perfused, fixed for 24 h in 4% PFA at 4°C, and embedded in paraffin (Leica ASP200S). 3-5 µm thick lung sections were prepared.

#### Preparation of lung tissue for paraffin sections

For paraffin embedding, the chest cavity was opened and the lungs were perfused with 25 ml HBSS, removed, and fixed in 4% PFA for 24 h at 4°C. Lungs were paraffin-embedded using (Leica ASP200S), sectioned at 3–5 μm thickness, and stained with hematoxylin and eosin (H&E) or immunofluorescence antibodies for histological analysis.

#### Staining of paraffin lung sections

For immunofluorescence or H&E-staining, lung sections were deparaffinized. For immunofluorescence staining, the sections were incubated in an antigen retrieval buffer (ThermoFisher) according to the manufacturer’s instructions. Following permeabilization (0.5% Triton-X-100 in PBS-/-) for 20 min and blocking (10% horse serum, 1% BSA in PBS-/-) for 1 h at 21°C, the sections were incubated with the following primary and secondary antibodies: rat anti-Ki67 IgG (14-5698-82, Invitrogen), Rabbit anti-ProSPC (ab90716, abcam), and goat anti-eCadherin (AF748-SP, R&D Systems) with goat anti-rat IgG AF555 (A-48263, Invitrogen), donkey anti-rabbit IgG AF647 (ab150075, abcam),and donkey anti-goat IgG AF488 (ab150129, abcam). Lung tissue sections were mounted with ProLong™ Diamond Antifade Mountant (ThermoFisher).

#### Preparation and immunofluorescence staining of vibratome sections

Precision-cut lung slices were generated as previously described^47^. Briefly, lungs were perfused, inflated with 2% (w/v) ultra-low melting point agarose in PBS (without Ca²⁺/Mg²⁺), and fixed in 4% paraformaldehyde for 24 h at 4 °C. Lobes were sectioned into 200–300 µm slices using a vibratome. For immunofluorescence staining, vibratome sections were incubated in blocking/permeabilization buffer (5% (w/v) BSA and 0.3% (v/v) Triton X-100 in PBS) for 1 h at room temperature. Primary antibodies diluted in blocking/permeabilization buffer were applied for 8 h at 4 °C. After washing with PBS, slices were incubated with secondary antibodies for 2 h at room temperature. The following antibodies were used: rabbit anti-ProSPC (ab90716, Abcam) and rat anti-RAGE (MAB1179, R&D Systems), followed by goat anti-rabbit IgG Alexa Fluor 647 (A21245, Invitrogen) and donkey anti-rat IgG Alexa Fluor 488 (A21208, Invitrogen). Slices were stored at 4 °C in PBS containing 0.05% sodium azide until imaging.

#### Immunofluorescence staining of murine organoids

The following primary antibodies were utilized: rabbit anti-mouse ProSPC (WRAB-9337, Seven Hills), mouse anti-human/mouse ProSPC (ab167608, Abcam), rabbit anti-mouse CAV-1 (ab2910, Abcam), rabbit anti-mouse phospho-ACC (Ser79) (PA5-17725, Invitrogen), and rabbit anti-mouse p-mTOR (Ser2448) (SAB5700327, Merck). As secondary antibodies, goat anti-mouse IgG AF647 (ab150115, Abcam), chicken anti-rat IgG AF647 (A-21472, Invitrogen), goat anti-rabbit IgG AF555 (A-32732, Invitrogen), and donkey anti-rabbit IgG AF488 (A-32790, Invitrogen) were applied. After the organoids were fixed with 4% PFA for 10-15 min, they were incubated with permeabilization/blocking buffer (5% serum and 0.5% Triton-X-100 in PBS-/-) overnight at 4°C, followed by overnight incubation with the primary antibodies diluted in permeabilization/blocking buffer. After washing (2% serum and 0.3% Triton-X-100 in PBS-/-), the samples were incubated overnight with the secondary antibodies, followed by another three washing steps. For nuclear staining, NucBlue™ Fixed Cell ReadyProbes™ Reagent (Thermo Fisher) was utilized. The samples were mounted with ProLong™ Diamond Antifade Mountant (ThermoFisher). To analyze the signal intensity of an organoid IF staining, the Matrigel® (Corning) was removed as previously described^48^.

#### Microscopes and image analysis

Bright-field images of organoids and H&E-stained lung sections were acquired on the EVOS^TM^ FL Auto Imaging System (Thermo Scientific). Image analysis was done in Fiji. For confocal imaging, the Leica SP8 CLSM system (Leica Microsystems, Germany) running on LAS X 3.5.7 software was utilized. The confocal images were acquired and analyzed a custom-made macros of Fiji software.

#### Statistics

Data are presented as mean ± SEM for normally distributed data. An unpaired Student’s t-test assessed significance between two groups, and ANOVA for three or more groups. Normality was checked using Anderson-Darling, D’Agostino, Shapiro-Wilk, and Kolmogorov-Smirnov tests. For non-normally distributed data, median and interquartile ranges are shown, with Mann-Whitney test for group comparison (GraphPad Prism 9). Significance was set at p<0.05 (*p<0.05; **p<0.01; ***p<0.005; ****p<0.0001).

## Data availability

The bulk RNA sequencing data generated in this study have been deposited in the NCBI Gene Expression Omnibus (GEO) under accession number GSE336615 and are publicly available.

## Acknowledgments

The work was supported by Deutsche Forschungsgemeinschaft (DFG, German Research Foundation) under Germanýs Excellence Strategy – EXC 2026, Cardio-Pulmonary Institute, Project ID: 390649896 to SH; SFB-TR84 (project number 114933180 to SH), SFB1021 (project number 197785619 to SH), and KFO309 (project number 284237345 to SH, REM); the German Center for Lung Research (DZL, project number 82DZL005B1 to SH); Institute of Lung Health (ILH) project number 82DZL005B4 to SH); the Hessen State Ministry of Higher Education, Research and the Arts (HMWK, Landes-Offensive zur Entwicklung Wissenschaftlich-ökonomischer Exzellenz, LOEWE, Förderlinie 4a project ID III L7– 519/05.00.002 to S.H.); the German Center for Infection Research (DZIF) partner site Giessen (TTU01.830_00, project number 8032801830 to SH); Federal Ministry of Education and Research (BMFTR, IPSELON 2.0 project number 03LW0521 to SH). EEA was funded by the Institute for Lung Health (ILH), DFG (EL 931/4-2, EL 931/5-1, EL 931/4-1, EL 931/2-2 (KFO309 284237345 Project P7), and SFB 1213 Project-ID 268555672 Project A04), the Excellence Cluster Cardio-Pulmonary System/Cardio-Pulmonary Institute (EXC 2026, project number 390649896), and DZL (82DZL005C1). AIVA was supported by Transdisciplinary Research Area Life and Health, Life & Medical Science Institute, excellence cluster ImmunoSensation, Project-ID 432325352 -SFB 1454 and SPP2493 (HetCCI).

## Author contributions

Conceptualization, SH, and AIVA; methodology, TS, AK, IA, MH, AIVA, and EEA; investigation, TS, AK, IA, MF, LS, LPO, AIVA, BO and TH; writing - original draft, SH, AIVA and TS; writing - review & editing, SH and AIVA; funding acquisition, SH; resources, IA, SH; supervision, SH and AIVA.

