## Supplementary Figures for "Granulocyte-macrophage colony stimulating factor targets lung stem cell niches to accelerate alveolar repair after virus-induced lung injury"

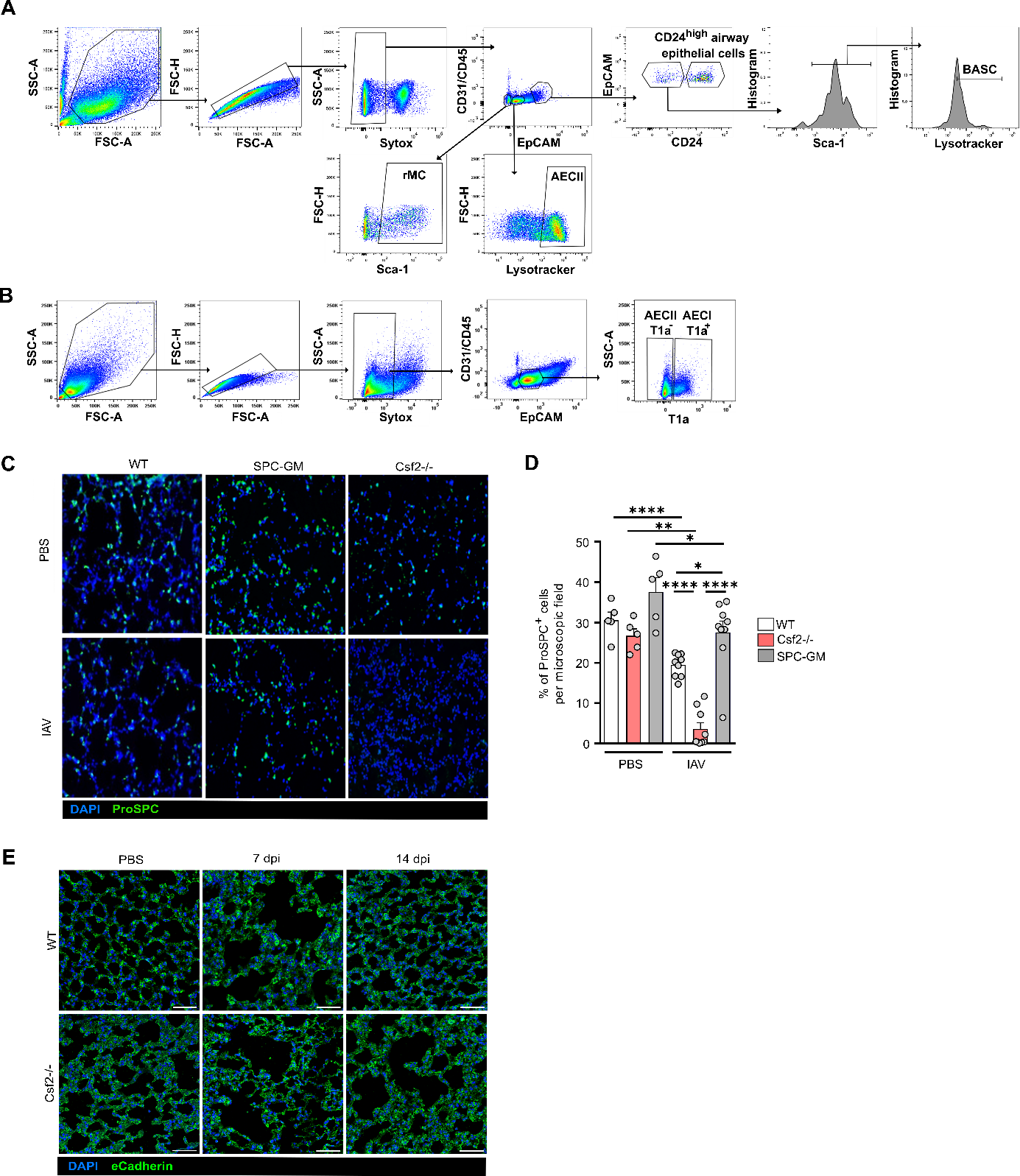

**Supplemental Figure 1. IAV infection enhances GM-CSF expression in alveolar epithelial stem cells increasing the amount of AECII and epithelial repair.** **(A)** Gating strategy for the isolation of living (Sytox-) AECII (CD31/CD45^neg^EpCAM^+^Lysotracker^+^), rMC (CD31/CD45^neg^EpCAM^neg^
Sca-1^+^), airway epithelial cells (CD31/CD45^neg^EpCAM^high^CD24^high^), and BASC (CD31/CD45^neg^
EpCAM^high^CD24^low^Sca1^+^Lysotracker^+^) from lung homogenate of infected WT or *Csf2^-/-^* mice. **(B)** Gating strategy for the isolation of living (Sytox-) AECs (CD31/CD45^neg^EpCAM^+^) type I (T1α^+^) and type II (T1α^-^) from lung homogenate of infected WT mice. **(C-D)** WT, *Csf2^-/-^*, and SPC-GM mice were infected with 500 FFU, and lungs were harvested at 7 dpi. The percentage of ProSPC^+^ cells (AECII and BASC) per high-power field was analyzed (n=5-9, two-way ANOVA followed by Tukey’s post hoc test). **(E)** Representative images of eCadherin-stained lung sections from WT (500 FFU) and *Csf2^-/-^* (100 FFU) mice at 7 and 14 dpi versus untreated controls (scale bar 150 µm). Graph shows means ± SEM; *=p<0.05; **=p<0.005; ****=p<0.0001.

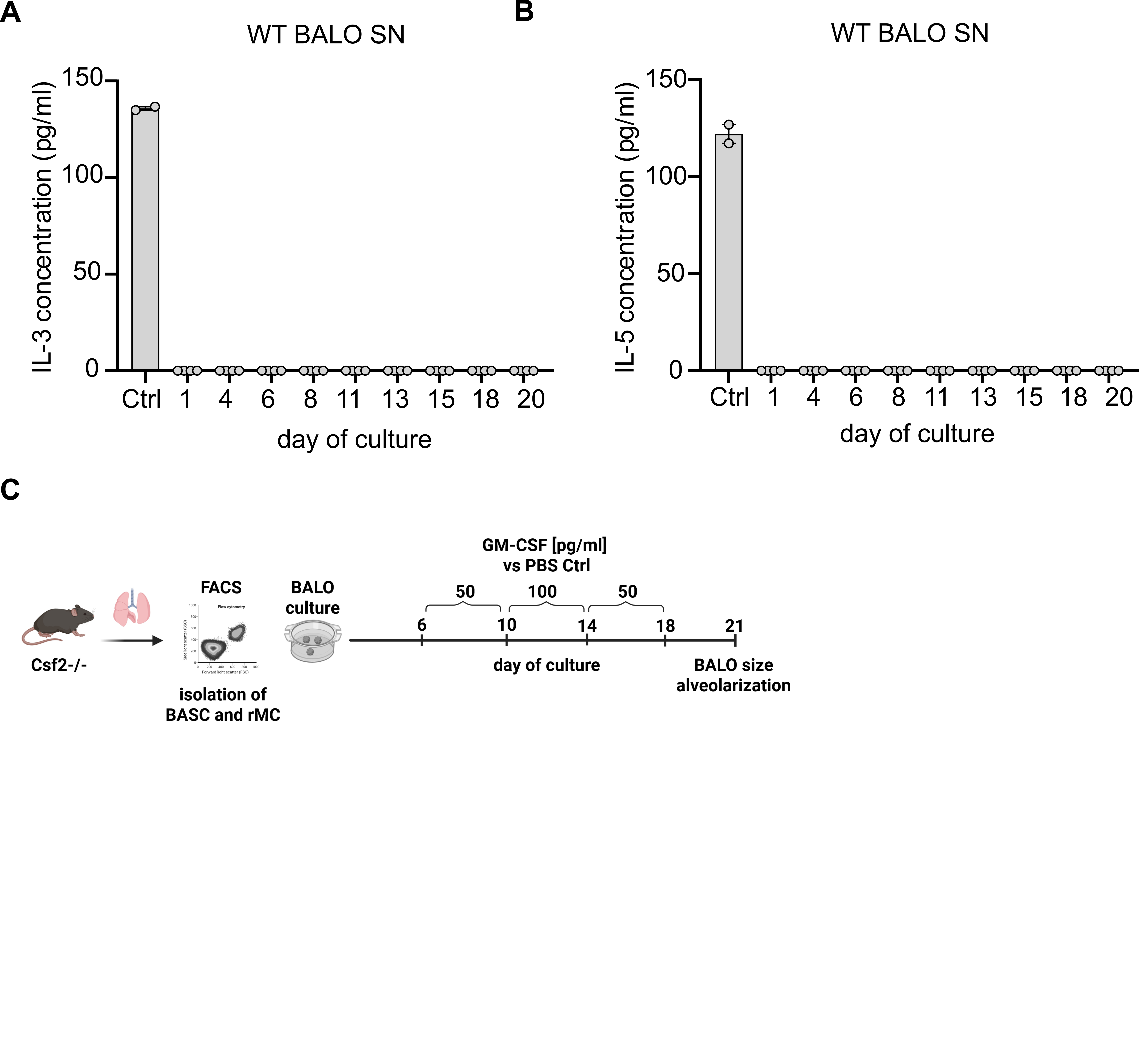
**Supplemental Figure 2. IL-3 and IL-5 is not secreted by WT BALOs while *ex vivo* rmGM-CSF stimulation promotes *Csf2^-/-^*** **BALOs development.** **(A)** IL-3 and **(B)** IL-5 concentrations (pg/ml) in WT BALO SN during culture (n=2-4). **(C)** Experimental setup: BASCs and rMCs were isolated from *Csf2^-/-^* mice for BALO co-culture. Exogenous recombinant GM-CSF was supplemented to the medium: 50 pg/ml from day 6 to 10, 100 pg/ml from day 10 to 14, and 50 pg/ml from day 14 to 21.

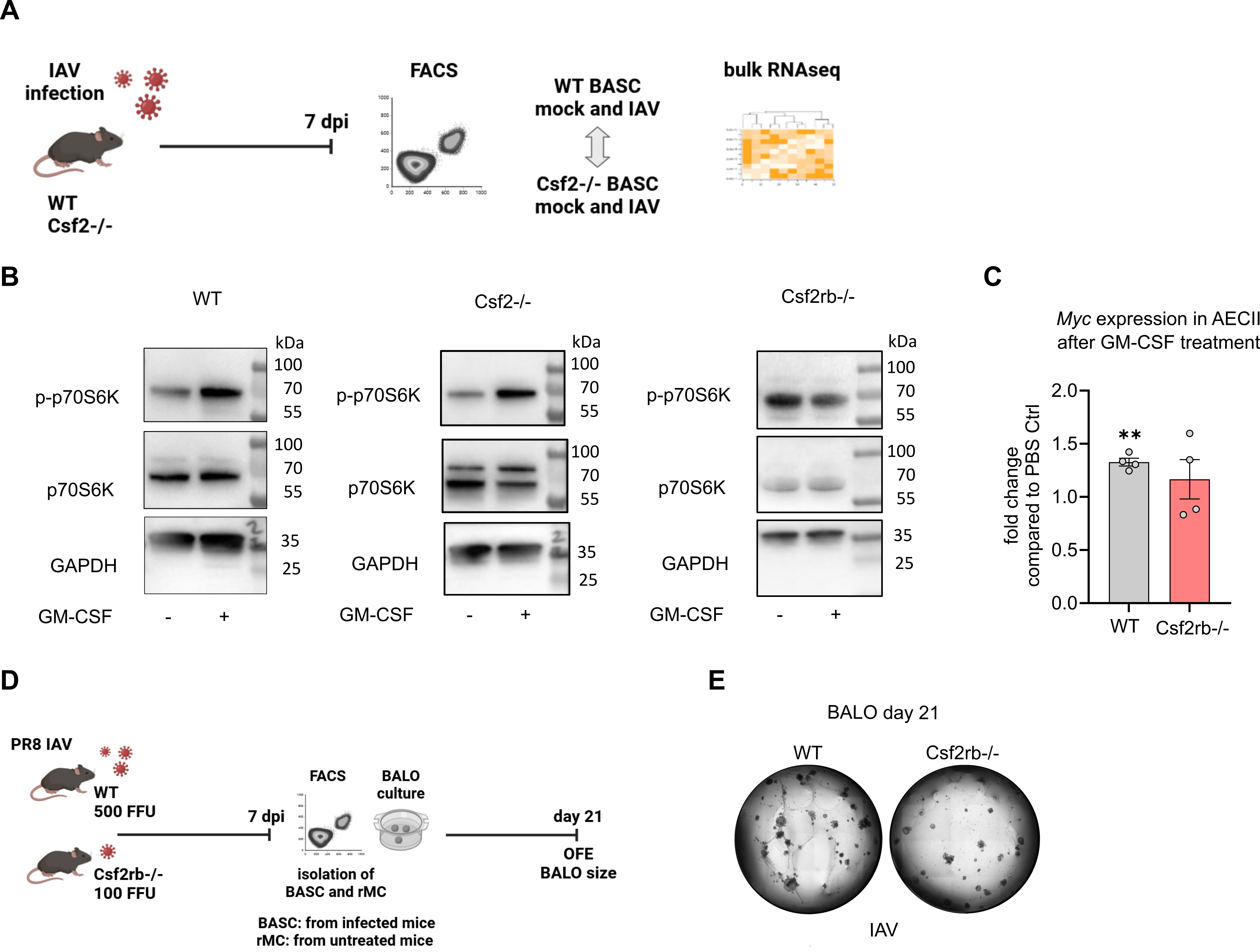
**Supplemental Figure 3. GM-CSF induces the expression of the cell proliferation in AECIIs *ex vivo*. (A)** Experimental setup: BASCs isolated from WT and *Csf2^-/-^* mice, IAV-infected (500 FFU, 7 dpi) or mock-treated, were subjected to bulk RNA-Sequencing. **(B)** Primary murine AECs from WT, *Csf2^-/-^,* and *Csf2rb^-/-^* mice were infected with PR8 IAV for 24 h *ex vivo* and treated with recombinant GM-CSF. Representative Western Blot images depict (p-)p70S6K (70 kDa) and GAPDH (37 kDa). **(C)** *c-Myc* expression in AECIIs that were treated with recombinant GM-CSF (50 ng/ml) *ex vivo* compared to PBS control (n=4, independent two-tailed Student’s t-test). Graph shows means ± SEM; **=p<0.005. **(D)** Experimental setup: BASC were isolated from WT and *Csf2rb^-/-^* at 7 dpi for BALO co-culture with rMC from untreated WT or *Csf2rb^-/-^* mice. **(E)** Representative transmission microscopy images of BALOs derived from WT (500 FFU) or *Csf2rb^-/-^* (100 FFU) BASCs of IAV-infected at 7 dpi.

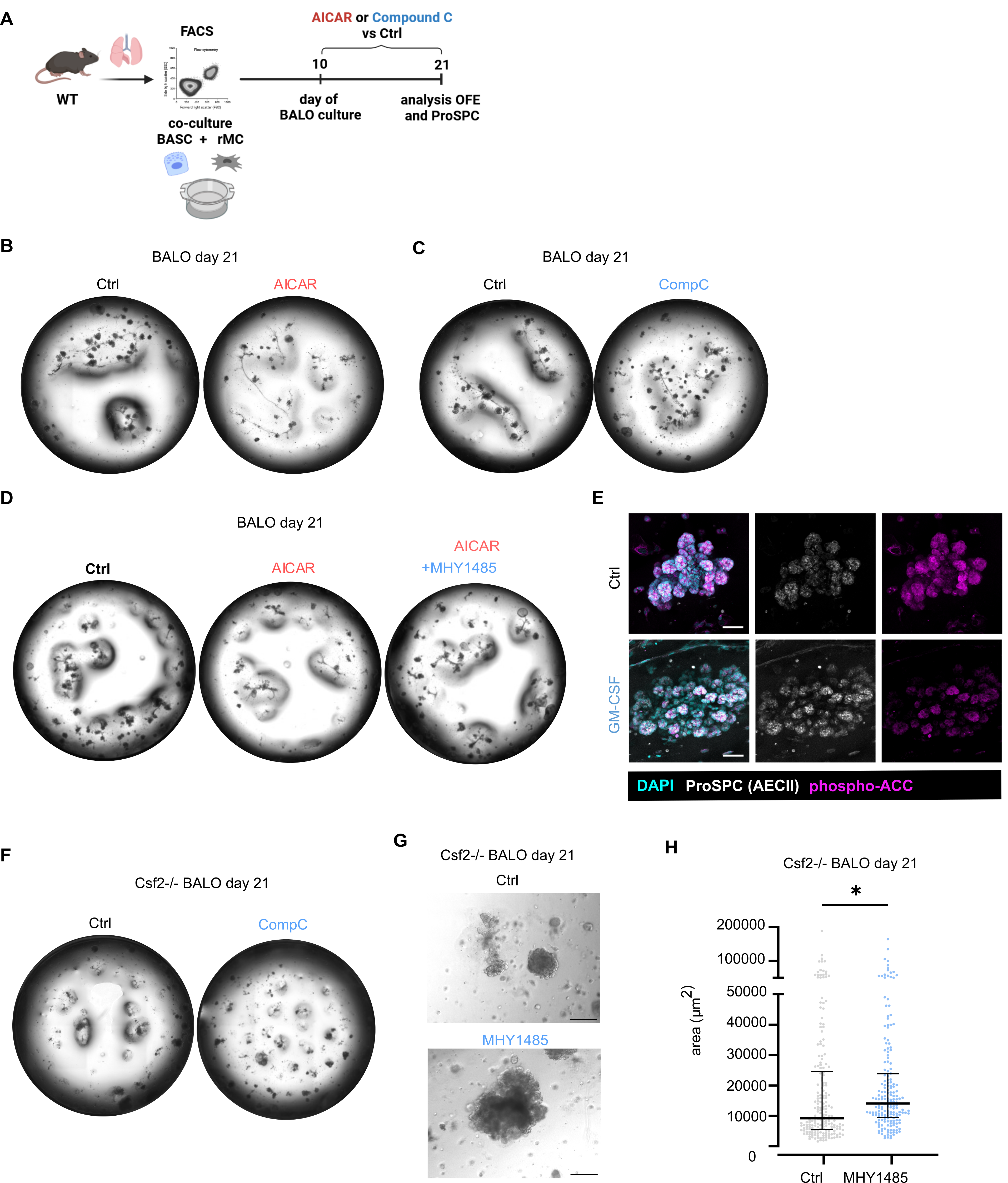

**Supplemental Figure 4.** **AMPK signaling modulation impacts organoid formation and alveolarization in BALOs.** **(A)** BALO cultures were generated by co-culturing FACS-isolated rMCs and BASCs from WT mice. On day 10, the treatment with AMPK activator AICAR (0.5 mM) or inhibitor Compound C (1 µM) was started. Organoid formation efficiency (OFE), ProSPC+ area, and number of alveoli were analyzed on day 21. **(B-C)** Representative transmission microscopy images of BALOs treated with **(B)** AICAR vs PBS or **(C)** Compound C vs DMSO. **(D)** Representative confocal images of ProSPC-stained BALOs at day 21. **(D)** Transmission microscopy of WT BALOs treated with AICAR (0.5 mM) or AICAR+MHY1485 (1 µM) vs control from day 10 to 21 of culture. **(E)** Confocal images of BALOs (day 12) treated with GM-CSF (50 ng/ml) vs PBS Ctrl from day 6 to 12 showing ProSPC and phospho-ACC (AMPK activity). **(F-G)** Representative transmission microscopy images of *Csf2^-/-^* BALOs treated with CompoundC or MHY1485 vs DMSO Ctrl. MHY1485 concentration: 0.5 µM (day 6-10), 1 µM (day 10-14), 0.5 µM (day 14-18). Compound C concentration: 0.5 µM (day 6-10), 1 µM (day 10-14), 0.5 µM (day 14-18). Scale bar 200 µm. **(H)** Size of *Csf2^-/-^* BALOs (µm²) after MHY1485 or DMSO Ctrl treatment (n=3-4 technical replicates, median with interquartile ranges are shown, Mann-Whitney test). *=p<0.05; **p<0.005; ****p<0.0001.

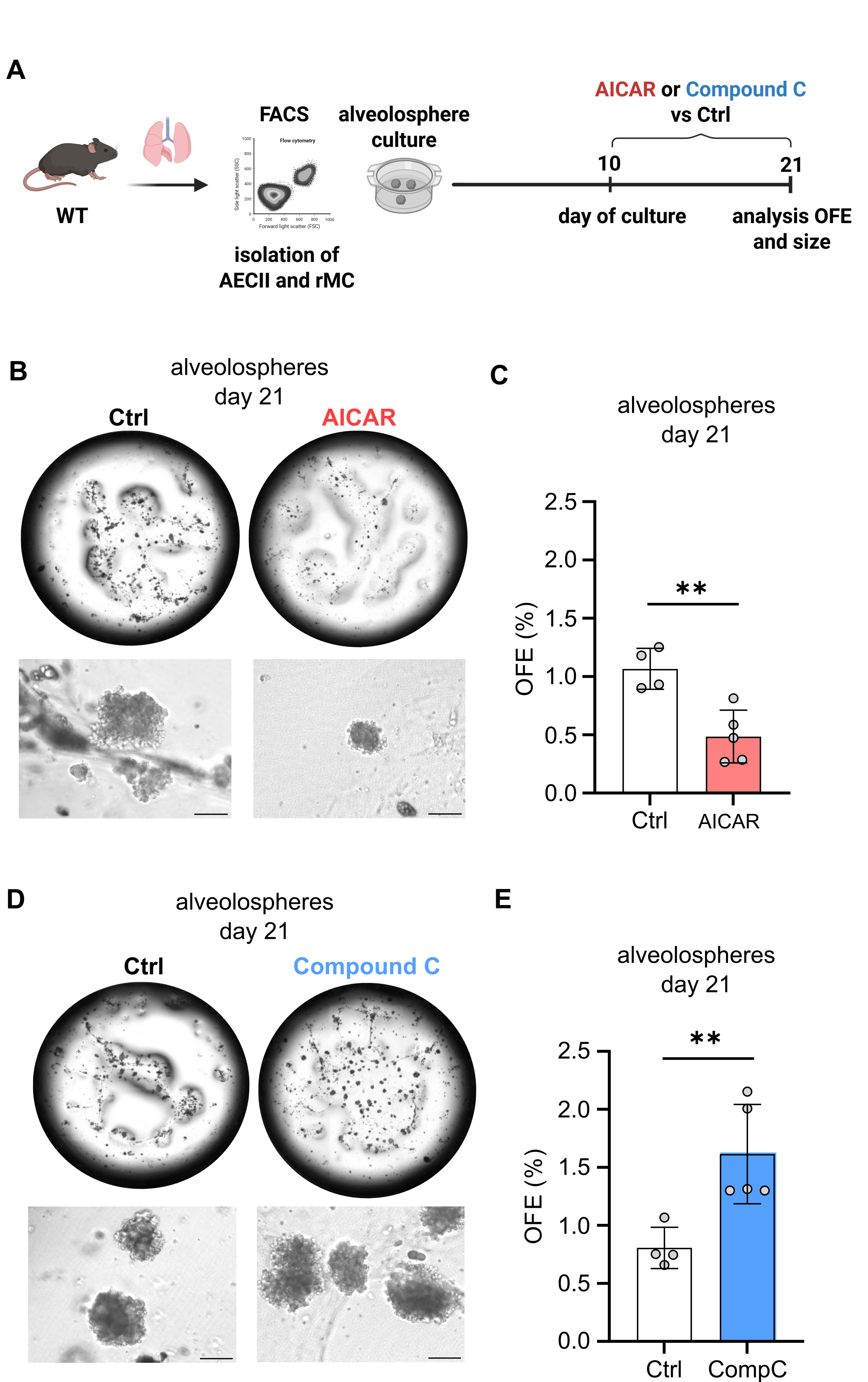

**Supplement Figure 5. AMPK inhibition increases whereas AMPK activation reduces alveolosphere growth. (A)** Scheme of experimental setup: AECs and rMCs were isolated from lung homogenate of adult WT mice and co-cultured for the generation of alveolospheres. Cultures were treated from day 10 to 21 with AMPK activator AICAR (0.5 mM) vs PBS Ctrl or AMPK inhibitor Compound C (1 µM) vs DMSO Ctrl. **(B, D)** Representative transmission microscopy images and **(C, E)** OFE (%) of alveolosphere cultures are shown (n=4-5 technical replicates, independent two-tailed Student’s t-test). Graphs show means ± SEM; **=p<0.005.

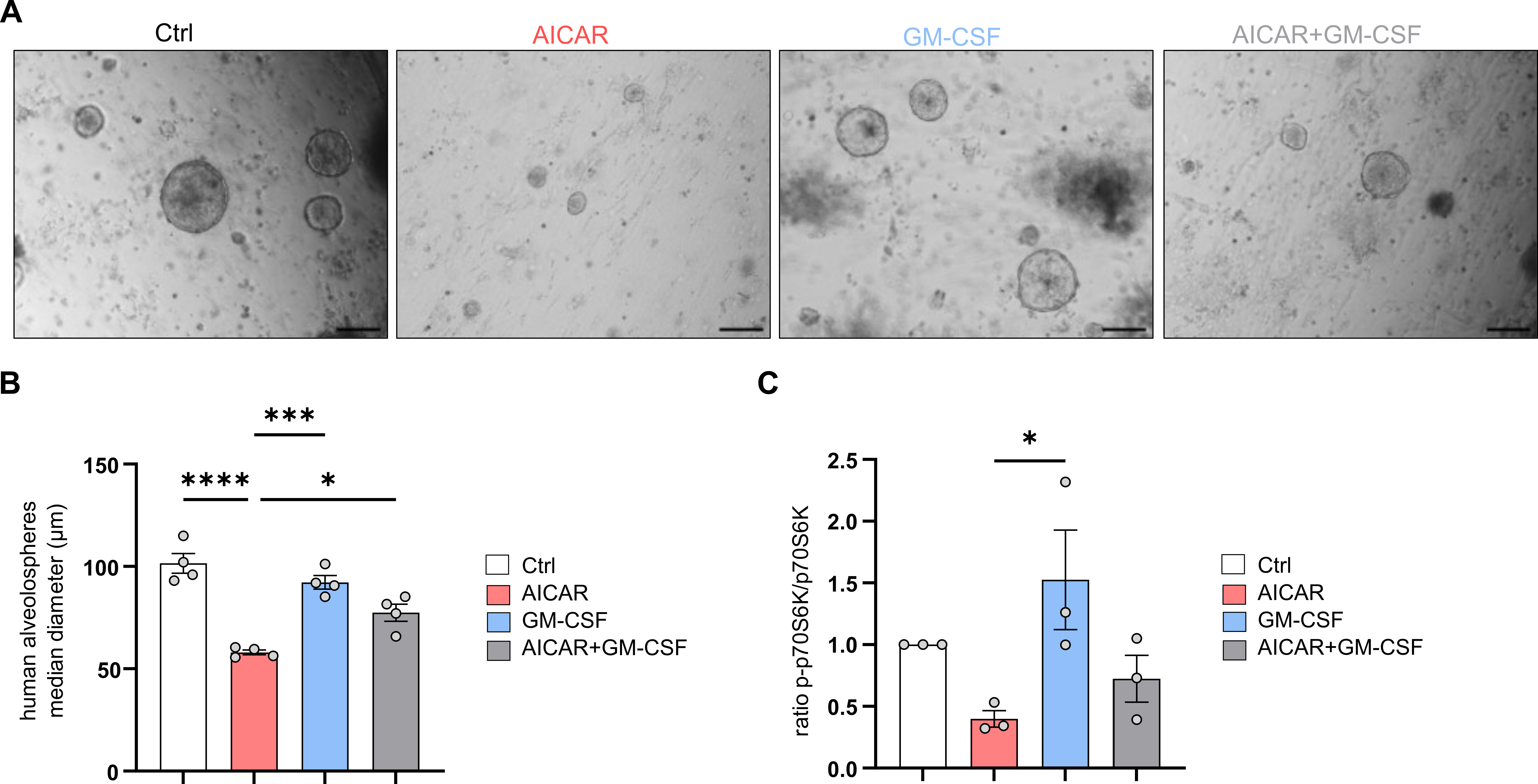

**Supplement Figure 6. Human alveolospheres growth is reduced by AMPK activator treatment but enhanced by GM-CSF-induced mTORC1 activation. (A)** Representative transmission microscopy images of human alveolospheres treated with AICAR (0.5 mM), GM-CSF (50 ng/ml), or AICAR+GM-CSF versus PBS control from day 28 to 35 of culture (scale bar 150 µm). **(B)** Analysis of the alveolospheres’ diameter (in µm) (n=4 technical replicates, one-way ANOVA followed by Tukey’s post hos test) and **(C)** the mTORC1 activity (phospho-p70S6K/p70S6K) analyzed in a Western Blot (n=3, one-way ANOVA followed by Tukey’s post hoc test). Graphs show mean ± SEM; *=p<0.05; ***=p<0.001; ****=p<0.0001.

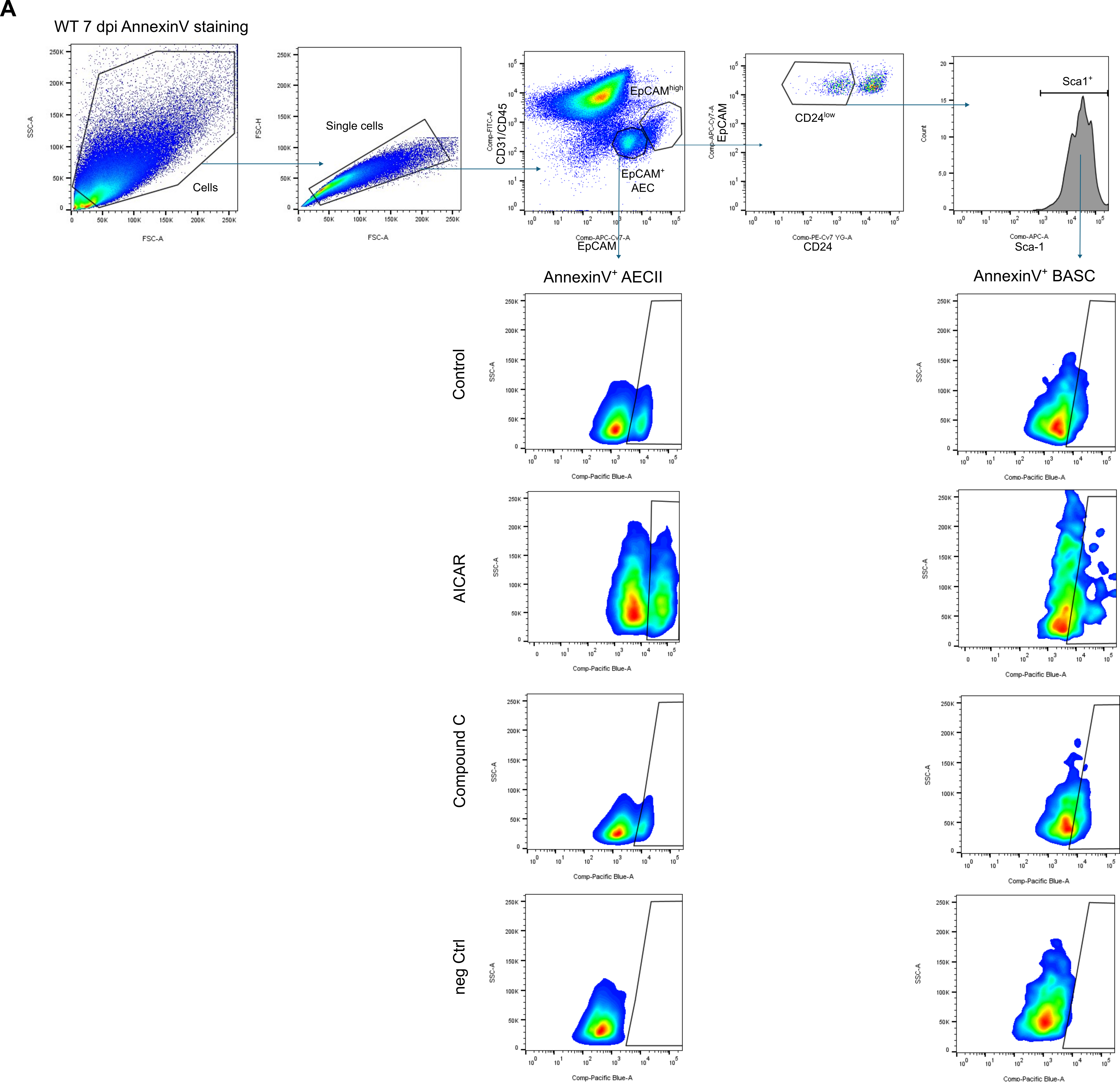

**Supplement Figure 7. AMPK signaling modulation affects AECII and BASC apoptosis *in vivo.*** Gating strategy for the analysis of AnnexinV^+^ AECIIs and BASCs. **(A)** AEC (CD32/CD45^-^EpCAM^+^) and BASC (CD32/CD45^-^EpCAM^high^CD24^low^Sca-1^+^) populations were analyzed for AnnexinV^+^ cells. Cell debris and cell doublets were excluded. Representative FACS plots are shown.

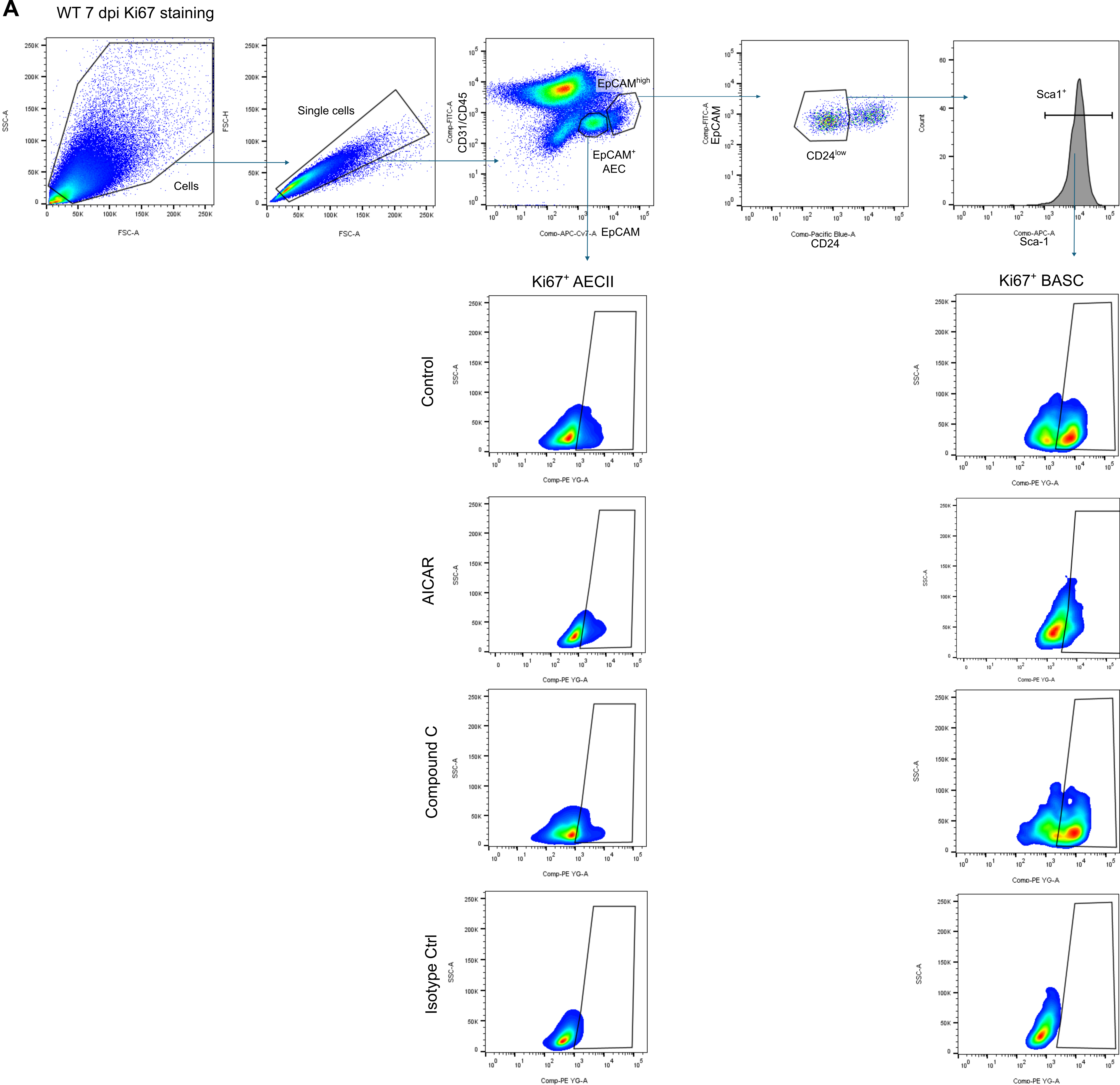

**Supplement Figure 8. AMPK signaling regulates proliferation of AECIIs and BASCs *in vivo*.** Gating strategy for the analysis of Ki67^+^ AECIIs and BASCs. **(A)** AEC (CD32/CD45^-^EpCAM^+^) and BASC (CD32/CD45^-^EpCAM^high^CD24^low^Sca-1^+^) populations were analyzed for the expression of Ki67. Cell debris and cell doublets were excluded. Representative FACS plots are shown.

**Supplement Figure 9. GM-CSF improves alveolar epithelial repair after IAV infection. (A)** Analysis of H&E-stained paraffin lung sections obtained from IAV-infected WT mice at 14 dpi that were treated with 10 µg GM-CSF vs PBS control at 3 and 5 dpi (scale bar 100 µm). **(B)** The vertical mean linear intercept (MLI in µm) was analyzed (n=3-4 mice, 4 images per mouse, independent two-tailed Student’s t-test). **(C)** Gating strategy for the analysis of tdTomato^+^ AECI (EpCAM^low^T1α^+^tdTomato^+^) in SPC-Cre^ERT2; tdTomato^ mice. Cell debris, doublets, and dead cells were excluded. **(D)** Representative confocal image of precision cut lung slice from SPC-Cre^ERT2,tdTomato^ mouse stained for ProSPC (AECII), RAGE (AECI), and DAPI with tdTomato^+^ cells. Zones 1 (healthy/mild damage), 2 (area of repair), and 3 (severe damage) are depicted. Scale bar upper panel 1 mm and lower panel 100 µm. Graph shows means ± SEM; *=p<0.05.
